# Speech decoding from a small set of spatially segregated minimally invasive intracranial EEG electrodes with a compact and interpretable neural network

**DOI:** 10.1101/2022.06.07.495084

**Authors:** Artur Petrosyan, Alexey Voskoboinikov, Dmitrii Sukhinin, Anna Makarova, Anastasia Skalnaya, Nastasia Arkhipova, Mikhail Sinkin, Alexei Ossadtchi

## Abstract

**Background:** Speech decoding, one of the most intriguing BCI applications, opens up plentiful opportunities from rehabilitation of patients to direct and seamless communication between human species. Typical solutions rely on invasive recordings with a large number of distributed electrodes implanted through craniotomy. Here we explored the possibility of creating speech prosthesis in a minimally invasive setting with a small number of spatially segregated intracranial electrodes.

**Methods:** We collected one hour of data (from two sessions) in two patients implanted with invasive electrodes. We then used only the contacts that pertained to a single sEEG shaft or an ECoG stripe to decode neural activity into 26 words and one silence class. We employed a compact convolutional network-based architecture whose spatial and temporal filter weights allow for a physiologically plausible interpretation.

**Results:** We achieved on average 55% accuracy using only 6 channels of data recorded with a single minimally invasive sEEG electrode in the first patient and 70% accuracy using only 8 channels of data recorded for a single ECoG strip in the second patient in classifying 26+1 overtly pronounced words. Our compact architecture did not require the use of pre-engineered features, learned fast and resulted in a stable, interpretable and physiologically meaningful decision rule successfully operating over a contiguous dataset collected during a different time interval than that used for training. Spatial characteristics of the pivotal neuronal populations corroborate with active and passive speech mapping results and exhibit the inverse space-frequency relationship characteristic of neural activity. Compared to other architectures our compact solution performed on par or better than those recently featured in neural speech decoding literature.

**Conclusions:** We showcase the possibility of building a speech prosthesis with a small number of electrodes and based on a compact feature engineering free decoder derived from a small amount of training data.

## 1 Introduction

Brain-computer interfaces (BCIs) directly link the nervous system to external devices [18] or even other brains [42]. While there exist many applications of BCIs [1], clinically relevant solutions are of primary interest since they hold promise to rehabilitate patients with sensory, motor, and cognitive disabilities [33],[13].

BCIs can deal with a variety of neural signals [40, 31] such as, for example, electroencephalographic (EEG) potentials sampled with electrodes located on the scalp [32], or neural activity recorded invasively with intracortical electrodes penetrating cortex [23] or placed directly onto the cortical surface [47]. A promising and minimally invasive way to directly access cortical activity is to use stereotactic EEG (sEEG) electrodes inserted via a burr hole made in the skull. Recent advances in implantation techniques including the use of brain’s 3D angiography, MRI and robot-assisted surgery help to further reduce the risks of such an implantation and make sEEG technology an ideal trade-off for BCI applications [21]. ECoG strips is another method to achieve direct electrical contact with cortical tissue with minimal discomfort to a patient [2].

The ability to communicate is vital to humans and speech is the most natural channel for it. Inability to speak dramatically affects the quality of life. A number of disorders can lead to a loss of this vital function, for example, cerebral palsy and stroke of the brain stem. Also, in some cases severe speech deficits may occur after a radical brain tissue removal surgery in oncology patients. While several technologies have been proposed to restore the communication function they primarily rely on brain controlled typing or imaginary handwriting [55] and appear to be practical only for severely affected patients. At the same time only in the United States 50 million people suffer from not being able to use their speech production machinery properly. A significant fraction of them have pathology not amenable by alaryngeal voice prosthesis [27] or “silent speech” devices [16] and require a neurally driven speech restoration solution.

Several successful attempts of BCI based speech restoration have already been made and a significant progress is achieved in decoding phonemes [56, 44, 38], individual words [34, 37, 51], continuous sentences [34, 37, 51] and even acoustic features [20, 51, 4] followed by the speech reconstruction algorithms using either Griffin-Lim or deep neural network algorithms inspired by WaveNet[4].

These solutions employ a broad variety of machine learning approaches for decoding speech from brain activity data. Starting from linear models [56], LDA [5], metric models [20] to deep neural networks (DNN) [34, 37, 51], that in general do not require manual feature engineering and can be applied directly to the data, however sometimes operating over a set of handcrafted features primarily derived from high-gamma activity. Several different neural network architectures have been tried for the speech decoding task: 1) relatively shallow ones consisting of a few convolutional or LSTM layers, 2) truly deep architectures with inception blocks [51] or with skip connections exploiting residual learning technique [4] as well as those borrowed from the computer vision applications [24, 52], 3) ensembles of DNN [37] making final solution more robust. Interestingly, that linear methods demonstrate compatible or, at least, close to DNNs decoding quality. Moreover, the latest studies obtained state of the art decoding accuracy using just a few layers over a set of handcrafted physiologically plausible features [34, 37]

The majority of the existing neural speech decoding studies rely on heavily multichannel brain activity measurements implemented with massive ECoG grids [37, 34, 4, 3] covering significant cortical area. These solutions for reading off brain activity are not intended for a long term use and are associated with significant risks to a patient [26] and suffer from a rapid loss of signal quality due to the leakage of the cerebrospinal fluid under the ECoG grid even if it is properly perforated. sEEG is a promising alternative whose implantation process is significantly less traumatic as compared to that of the large ECoG grids. The use of sEEG has already being explored for the speech decoding task [5] but the reported decoder again relied on a high count of channels from multiple sEEG shafts distributed over a large part of the left frontal and left superior temporal lobes which reduces the practicality of the proposed solution. A solution capable of decoding speech from the locally sampled brain activity would be an important step towards creating a speech prosthesis device.

The accuracy of neural speech decoding improves with the use of compressed representations encoding speech kinematic or acoustic features as an intermediate representation of the target variable [4] or for regularization [51]. However it still remains unclear which of the compressed speech representations is optimal for decoding speech from electrophysiological data and how it should be used to yield the best decoding accuracy. In addition to the direct practical benefit, answering this question together with an appropriate interpretation of the decision rule will shed light onto the neuronal basis and cortical representation of the speech production processes.

Here we explore the possibility of decoding individual words from intracranially recorded brain activity sampled with compact probes whose implantation did not require a full blown craniotomy. Our study comprises two subjects implanted either with sEEG shafts or ECoG stripes both via compact drill holes. We decode individual words using either 6 channels of data recorded with a single sEEG shaft or the 8 channels sampled using a single ECoG strip. For decoding we employed our compact and interpetable CNN architecture [43] augmented with the bidirectional LSTM layer [22] to compactly model local temporal dependencies in the internal speech representation that we used as the intermediate decoding target. We also compared the ultimate word decoding accuracy achieved with different internal representations. Our decoder operated causally using only the data from time intervals preceding the decoded time moment and therefore is fully applicable in a real-time decoding setting. Overall our study is the first attempt to achieve acceptable individual words decoding accuracy from cortical activity sampled with the compact non-intracortical probes whose implantation is likely to cause minimal discomfort to a patient and can be done even with local anesthesia.

## 2 Data

In this study we used two datasets collected from two epilepsy patients undergoing planned sEEG and ECoG implantation for the needs of presurgical mapping. The first patient was implanted bilaterally with a total of 5 sEEG shafts with 6 contacts in each with the goal to localize seizure onset zone. The implantation was performed under general anesthesia via five 3-mm drill holes. The second patient was implanted with 9 ECoG stripes of 8 contacts each covering frontal and inferior temporal lobes. The implantation was performed via several 12 mm drill holes. Figure 1 demonstrates post-surgical CT scans of the two patients. On the second day past the implantation both patients went through the active and passive [48] speech mapping procedures that yielded concordant results. In Patient 1 electrical stimulation of the 10-11 pair (300 *μs*, 2.5 mA, 50 Hz) resulted in pronounced speech arrest. The passive speech mapping procedure based on computing the mutual information (MI) between the speech envelope and the envelope of the gamma-band (60 Hz −100 Hz) filtered sEEG activity resulted into a sharp peak of the MI values for electrodes 9-12, see Figure 1.a,c. No speech related artefacts were observed when stimulating contacts 11-12 which could be due to the very sparing stimulation settings used in this patient - our stimulation current in this patient never exceeded 3 mV which is below the traditionally average current magnitude typically used for speech mapping [14]. Stimulation based speech mapping in Patient 2 caused involuntary tongue retraction when applied between electrodes 15-16 and the MI profile highlighted bluecontacts 13-15, see Figure 1.b,d.

**Figure 1:**
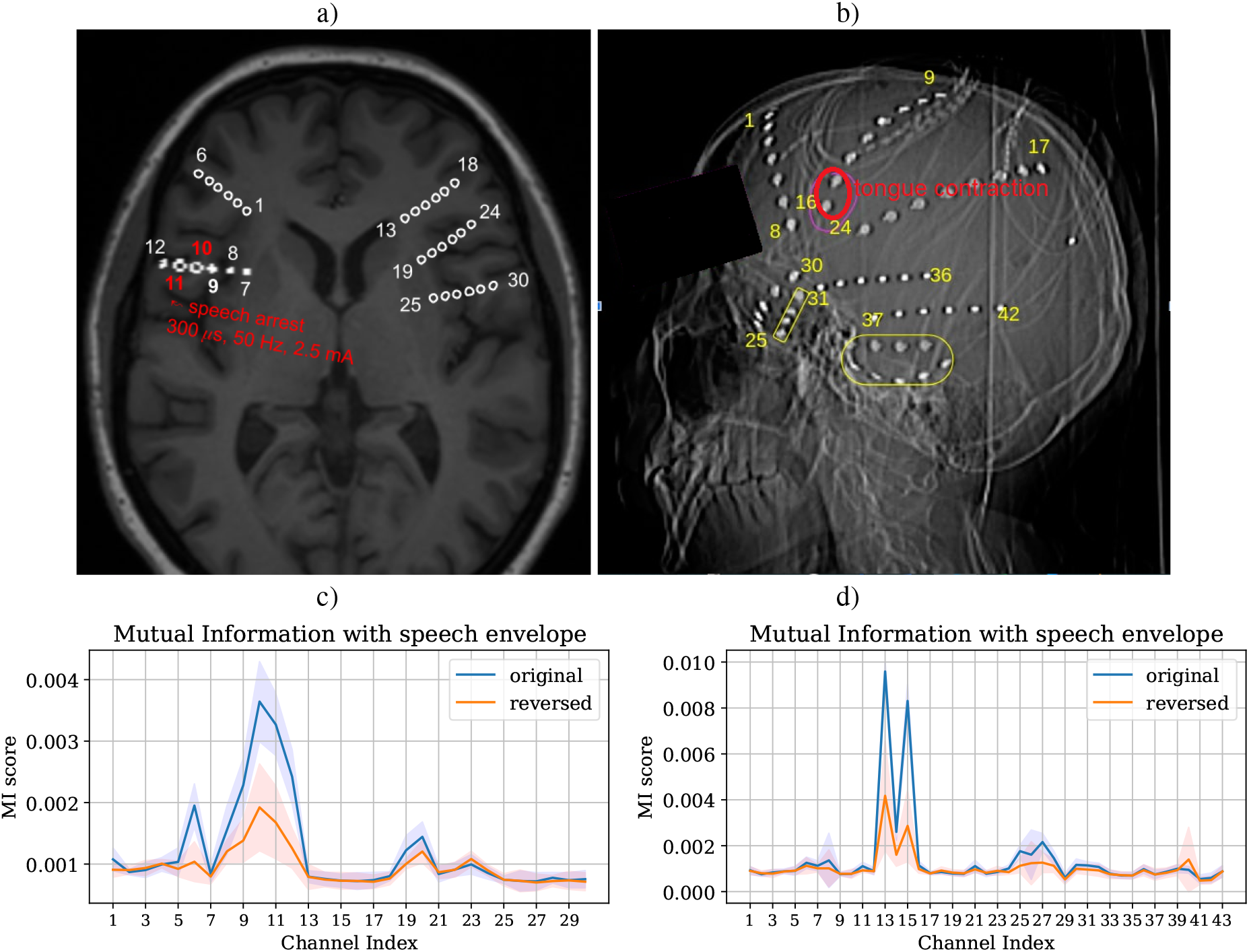
a) sEEG contacts extracted from the post-implantation CT scan of the first patient superimposed over her MRI. Bipolar electrical stimulation of the 10-11 pair (300 ts, 2.5 mA, 50 Hz) resulted in reproducible speech arrest. b) CT of the second patient who was implanted with nine 8-contacts ECoG stripes covering bilateral frontal and inferior temporal lobes. Bipolar electrical stimulation applied to electrodes 15-16 caused involuntary tongue retraction. c) Patient 1, mutual information profile between the speech envelope and gamma-band (60 Hz −100 Hz) filtered sEEG activity shows into a sharp peak of the MI values for electrodes 10 and 11. d) Patient 2, mutual information profile shows peak over electrodes 13-15 and in several other locations of this ECoG strip. The MI profiles between the time-reversed audio stream and original ECoG data is shown in red. The shadow corresponds to the standard deviation of the MI values estimated using the collection of different 3 minute long segments. The remaining bumps in the time-reversed MI profile may be due to the inherently rhythmic nature of the audio stream produced by the patient in response to the sequence of computer instructions.

Note that the exact shape of the MI profiles depends on the filtering parameters and therefore these plots need to be interpreted carefully. The MI profiles may also confuse speech production and perceptions processes, especially given the observation demonstrated in [30] that gamma activity in the auditory cortex accurately tracks the perceived speech envelope and may contribute to the observed MI.

The study was conducted according to the ethical standards of the 1964 Declaration of Helsinki. The participants provided written informed consent prior to the experiments. The ethics research committee of the National Research University, The Higher School of Economics approved the experimental protocol of this study. After the patients signed the appropriate informed consent we asked them to read off the succession of the 6 sentences presented at a comfortable pace in randomized order on the computer screen. Each sentence was repeated 30 and 65 times by the first and the second patient respectively. The sentences contained on average 4.3 words. Half of the sentences had direct and the other half indirect order of words and the majority of words within a single sentence started from the same letter. This was done to enable subsequent neurolinguistic analysis of the collected datasets. The sEEG in Patient 1 was recorded with an 80 channel g.HIamp amplifier. Patient’s 2 ECoG was registered with a 64 channel EBNeuro BE Plus LTM device. The sampling rate was set to *F_s_* =19200 Hz (Patient 1) and *F_s_* =4096 Hz (Patient 2). In both cases synchronously with neural activity we recorded speech signal measured with the Behringer XM8500 microphone.

## 3 Methods

### 3.1 Data preprocessing

We first parsed audio data into separate words. To this end, we manually processed several example word alignments and then used them to find similar ones by means of the dynamic time warping (DTW) algorithm [8]. Manual check shows that the absolute majority of the word alignments were detected correctly. For each word this procedure resulted into a list of index pairs corresponding to the start and the end of the word’s utterance. Audio data were processed using librosa Python software package [36] in order to extract log-mel spectral coefficients (LMSC) [50], mel-frequency cepstral coefficients (MFCC) [57] and several derivatives of the linear predictive coding (LPC) coefficients as described below. The sequences of these internal speech representations (ISRs) were downsampled to 1 KHz. We have experimented with all listed ISRs, see section 3.3 and Figures 7, 11.b in section 4. In the majority of the reported results we used LMSCs as the internal speech representation.

sEEG and ECoG data went through minimal pre-processing such as causal band-pass FIR filter in the 5-150 Hz frequency range. Then the data were resampled to 1 kHz sampling rate and the amplitude in each channel was standardized by subtracting the mean and dividing by the standard deviation. This multichannel data were used as an input of our decoding algorithm.

### 3.2 Decoding

In accord with the view expressed in [37] we decided to explore decoding accuracy on the level of individual words. On the one hand, words represent a sufficiently low level of detail which permits extension of the obtained solution into a broader range of application scenarios. On the other hand, words are less volatile as compared to phonemes as the articulation of the latter greatly varies depending on the flanker sounds neighboring the phoneme. This may mean that the neural encoding governing the transition between the different states of the articulatory tract may vary significantly from case to case depending on the phonetic context a phoneme is encountered in.

### 3.3 Internal speech representations

Most of the ISRs are based on modeling speech signal as produced by an excitation sequence passing through a linear time-varying filter [25]. The excitation sequence is the air flow in the larynx and the filter is formed by the articulatory tract elements (pharynx, vocal folds, tongue, lips, teeth) whose mutual geometry changes over time.

Linear predictive coding (LPC) and cepstral analysis are the two principal ways to estimate parameters of such a filter. LPC analysis is based on a direct estimate of the auto-regressive prediction coefficients (PC) *a_i_* through Burgs method [35]. However, prediction coefficients themselves are unstable, as their small changes may lead to large variations in the spectrum and possibly unstable filters. In order to decrease such an instability the following several equivalent representations are commonly used.

Reflection coefficients (RC) *k_i_* can be computed alongside with prediction coefficients through Burgs method and represent the ratio of the amplitudes of the acoustic wave reflected by and the wave passed through a discontinuity.

Another descriptor, log-area ratio (LAR) coefficients, *g_i_*, are equal to the natural logarithm of the ratio of the areas of adjacent sections in a lossless tube equivalent of the vocal tract having the same transfer function and can be computed from the reflection coefficients as 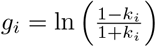.

Line spectral frequencies (LSF) is another highly efficient speech data compression technique [49] as errors in representing one coefficient generally result in a spectral change only around that frequency.

In what follows we will present the results of our experiments with several ISRs but our final decoding accuracy results are based on the use of log-mel spectral coefficients (LMSC).

#### 3.3.1 Synchronous decoding

Our goal is to decode specific words from the immediately preceding chunks of neural activity data. The direct approach would require gathering a large amount of training data. Instead we developed our decoding solution based on the idea described in [34] where the vocoder-like compact internal speech representation (ISR) is used for regularization purposes during the training. However, here instead of using the ISR as a regularizer we employ it as the intermediate target. In other words, we first use our LSTM-extended compact and interpretable architecture [43] to decode *M* = 40 log-mel spectral coefficients (LMSC) from either sEEG or ECoG based measurements of brain activity. After having trained this LMSC decoder optimizing the average correlation coefficient between the actual and the decoded LMSCs we fix its weights and train a simple convolutional neural network to decode discrete words based on the internal representations that emerged in the LMSC decoder. After training, our two-stage architecture operates as a single network on the minimally neural activity data and yields discrete classification of individual words at its output. Importantly, our experiments show that the use of the internal representations that emerged in the LSTM layer instead of the actual decoded LMSCs noticeably improves the final word classification accuracy. This observation is inline with a similar finding in a completely different domain [17] where the authors advocated the use of multiple separate “views” generated by different networks as the input to the final classifier in the image classification task.

For the training word classification task we manually extracted alignment of each word. We used only a chunk of neural data that corresponds to the particular word‘s alignment (we do not use information of neighborhood words). We also added a “silent” class that corresponds to the intervals of silence between word utterances. To get meaningful accuracy metrics we randomly drop a fraction of “silent” class samples to make the dataset class-balanced.

In this paradigm decoding of ISRs from neural data was performed asynchronously, i.e. on a rolling basis for each time point. In the causal regime, each time point was decoded based on the preceding 1000 ms of neural activity data, in the anti-causal mode we used 1000 ms window from the immediate future and in the non-causal mode we exploited two 1000 ms on both sides of the decoded time-point. The individual words decoding task was then accomplished synchronously, i.e. based on the decoded speech representations cut in the vicinity of each actual utterance.

#### 3.3.2 Asynchronous decoding

We have also experimented with a completely asynchronous approach illustrated in Figure 2. In contrast to the synchronous mode, in the asynchronous BCI setting our general task is to predict the uttered word based on the neural activity data preceding the word to be uttered (or the silence interval) at every time moment t. Hypothetically this information can then be used for speech generation.

**Figure 2:**
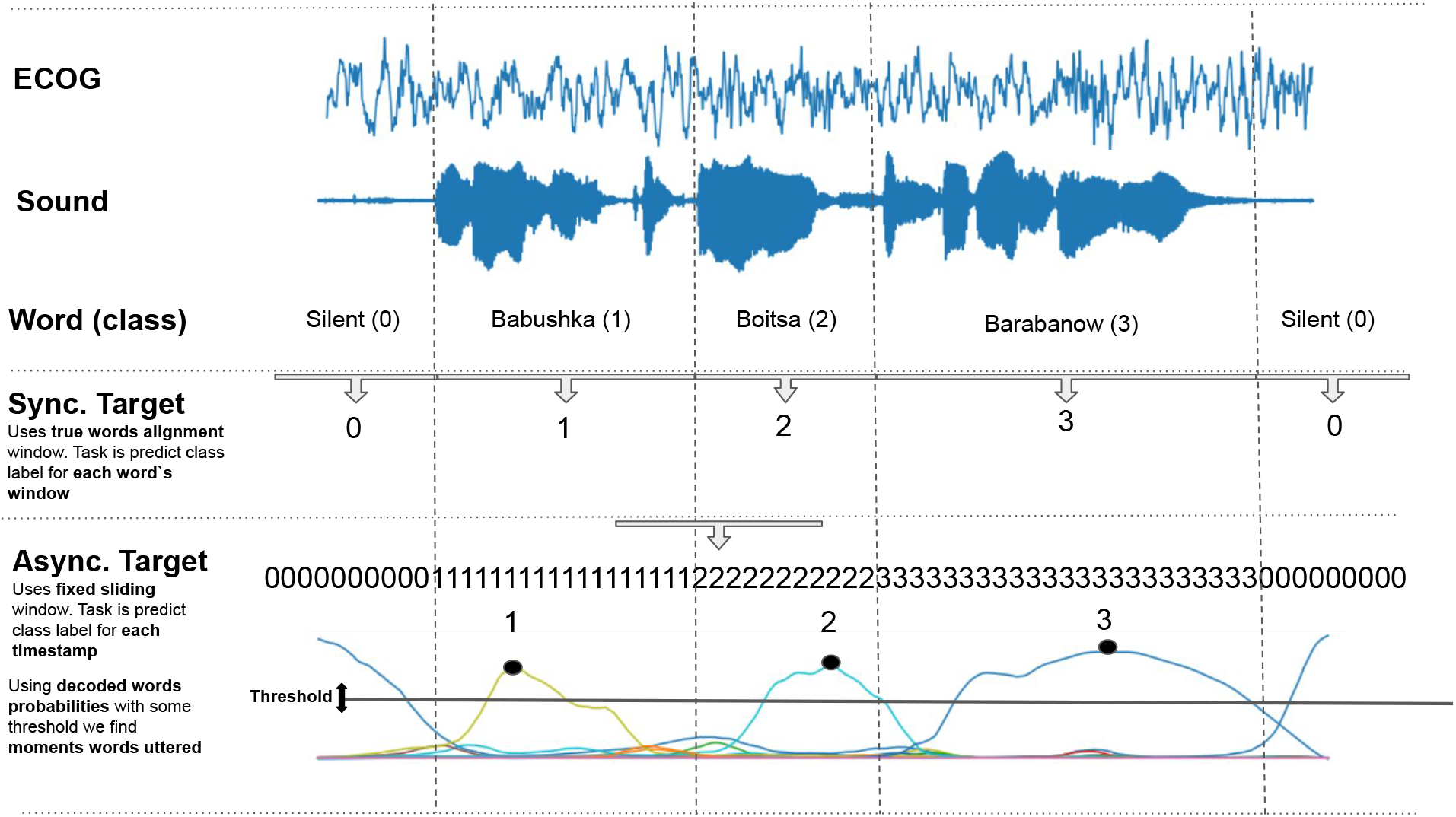
Illustration of the synchronous vs. asynchronous operation mode. In contrast to the synchronous mode, in the asynchronous regime our general task is to predict the uttered word based on the neural activity data for every time moment *t*.

Our first task here is to infer the probabilities *p_i_*(*t*) for each *i*–th word + the silence class for each time instance *t* based on the neural activity data [**x**(*t* – *T*, … **x**(*t*)] from the preceding time window of length *T*. As the next step we smooth the obtained probability profiles with 0.2 second moving average and choose the word (or the silence) based on the thresholding the smoothed probability profile 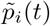. If 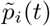 peaks and exceeds the threshold we make the corresponding decision and “utter” the *i*–th word. This very word can not be uttered again unless 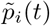 drops below, crosses the threshold and peaks again. See also section 3.5 and Figure 13 a) for additional clarifications.

### 3.4 Network architecture

For neural signals to LMSC decoding we employed the compact and interpretable convolutional network architecture developed earlier for motor BCI purposes [43] and augmented it with a single bidirectional LSTM layer with 30 hidden units to compactly model temporal regularities. The LSTM layer is followed by the fully connected layer with *M* = 40 output neurons each corresponding to a single mel-spectral coefficient whose temporal profile we are aiming to reconstruct from the neural activity data, see Figure 3. Note that unlike [5] we do not specify upfront the feature extraction parameters and let our architecture learn them during the training process guided by the optimization of the mean (across all MSCs) Pearson’s correlation coefficient between the original and the decoded LMSC timeseries. This feature extraction is performed by the adaptive envelope detector (ED) block that comprises a succession of the factorized spatial and temporal convolution operations followed by the rectification and smoothing blocks. The ED during training can potentially adapt to extracting instantaneous power of specific neuronal populations activity pivotal for the downstream task of predicting the LMSCs. In the search for the optimum, the ED weights are not only tuned to such a target source but also tune away from the interfering sources [19, 43]. The proper interpretation of the learnt ED’s weights allows for subsequent discovery of the target source geometric and dynamical properties.

**Figure 3:**
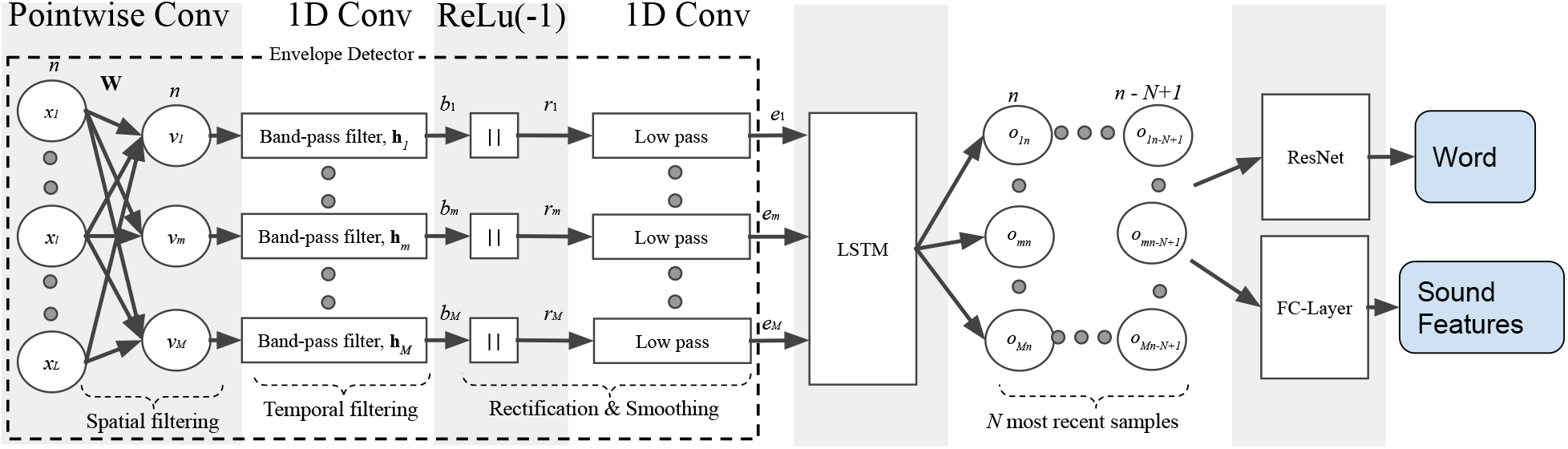
The architecture based on [43] and adapted for speech classification task. We used the same envelope detector technique to extract robust and meaningful features from the neuronal data. We then used the LSTM layer to account for the sequential structure of the speech ISR (e.g. LMSC) and finally decoded it with a fully connected layer over the LSTM hidden state (*o_ij_* on the figure). A separate 2D convolutional network was trained and used to classify separate words from the activity of thus pretrained LSTM.

**Figure 4:**
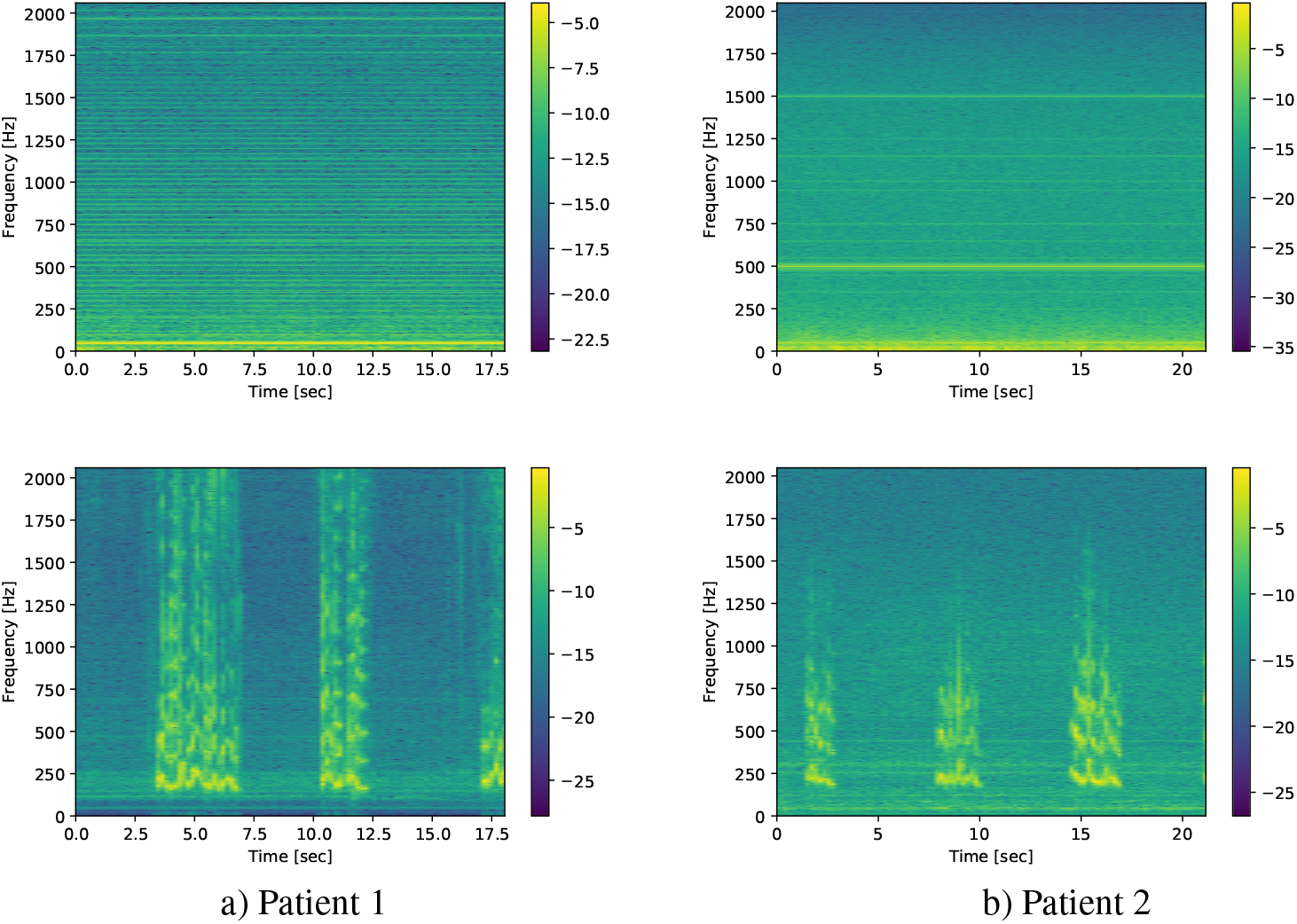
Log-spectrograms for audio and neural data in the electrode with the strongest audio-neural correlations

In order to measure the quality, we used 6 fold cross validation. In each of the folds we took 5/6 of independent sessions and 1/6 for validation, corresponding to 50 minutes and 10 minutes respectively. We used Adam optimiser for training with *α* = 0.0003 learning rate parameter. For training we used the entire train portion of our data, not just speech segments. Using only speech intervals worsened the quality of decoding by 15-20%. Our intuition here is that non-speech segments are also useful to the training and may serve regularization purposes. Also, hypothetically, the brain activity that determines the upcoming utterance happens during the silence interval and therefore not including this segment into the training could have detrimental effect on the final classification accuracy.

As described in section 4 in a number of our experiments we have replaced LMSCs with several other ISRs, see section 3.3, as a target for our first network.

After having trained our compact architecture to decode the ISRs as our intermediate target we used a 2D-convolution network to perform discrete classification of 26 words and the silent class using the representations developed in the one before the last layer of the compact architecture, see Figure 3.

### 3.5 Performance metrics

We use correlation coefficient to measure LMSC–from–neural activity reconstruction quality. To assess the words decoding quality when operating in the synchronous mode we downsample “silence” to avoid the positive bias in the reported numbers and then measure accuracy as the fraction of correctly classified utterances. We report our results in the form of 27 × 27 confusion matrices illustrating the proportion of correct and erroneous decoding of 26 words and the silence class.

To assess the accuracy when operating in the asynchronous mode we use precision-recall characteristics. As described earlier for each *i*–th word we compute smoothed probability profiles 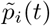 for each time instance *t*. We make a decision about a word being pronounced only at time points corresponding to the local maximums of 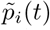 that cross the threshold *θ*. The *i–th* word is decoded if the local maximum of 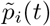 located above *θ* also appears to be the largest among all other profiles, i.e. 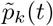, *k* ≠ *i*. In case the chosen *i*–th word corresponds to the one that is currently being uttered we mark this event as true positive (TP).

If after such a detection 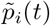 remains above the threshold and exhibits another local maximum which exceeds the values of all other smoothed probability profiles we will also make a decision to “utter” the *i*–th word. However, in this case this decision will be marked as false positive (FP) even if *t* belongs to the time range corresponding to the actual *i*–th word, because this results in the duplicated uttering and adds errors to the decoded words sequence. We also mark as FP the events when the index of the detected word does not match that of the actual pronounced word, see Figure 13 a) for the graphical representation of the above description. To compute these P-R curves we first smooth the probability profiles delivered by the neural network with a simple box-car averaging over the 0.2 sec segment. Then, we vary the detection threshold (single value for the entire test data segment) and compute the corresponding precision-recall pair. Doing so for a dense grid of thresholds we obtain a threshold independent metrics of algorithms performance.

We represent our asynchronous decoding results in the form of precision-recall curves parameterised by the threshold *θ* applied to probability profiles. Then, we use standard expressions and for each value of *θ* evaluate the precision and recall indicators as

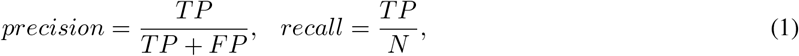

where *N* is the total number of utterances.

### 3.6 Weights interpretation

When dealing with overt speech decoding from neural activity data one needs to make sure that the obtained decision rule indeed uses neural activity data and does not exploit for decoding the possible artifacts such as electrical currents accompanying muscular activity or the acoustic signal leaked into neural data channels via, for example, microphone effect [45]. Thankfully, the widely spread over cortex spatial patterns of muscular activity occupying high frequency range [15] can be delineated from neural signals whose high frequency components, on the contrary, tend to be restricted to spatially compact cortical regions [53, 39]. To do so one needs access to both spatial and frequency domain patterns of the activity that appears pivotal to the decoder. Interpretable decision rules facilitate such tests for physiological plausibility of the obtained solutions. By extracting spatial and frequency domain patterns from the weights of the corresponding layers [19, 43] we can check for the physiological plausibility using domain specific knowledge as described above.

In this work we use our compact convolutional network as the front-end which allows for the theoretically justified interpretation of its spatial and temporal convolution weights by extracting spatial and frequency domain patterns corresponding to the neuronal populations pivotal to the specific downstream task. The details of our approach are outlined in [43] next we briefly review the basic ideas behind it.

The front-end of our network comprises factorized spatial and temporal convolution layers, see Figure 3. During training, the spatial and temporal filter weights of each branch not only get tuned to the pivotal neuronal sources but also tune away from the interfering signals.

In terms of spatial processing, that is combining the data from different sensors with specific weights, each branch of our adaptive envelope detector (ED), see Figure 3, corresponds to the model studied in [19]. However, each branch of the ED contains both spatial and temporal filters. Therefore, as we show in [43], the interpretation of branch’s *spatial* weights needs to be conducted withing the context set by the corresponding *temporal* filter. Since both spatial and temporal filtering are linear, interchanging them in the above statement is also valid and thus brunch’s temporal filter weights interpretation needs to be done taking into account the spatial filter of this branch. More formally our approach is summarized below.

Our ED processes data in chunks of a prespecified length of *N* samples. First, assume that the input segment length is equal to the filter length in the 1-D temporal convolution layer. Consider a chunk of input data from *L* channels observed over the interval of *N* time moments that can be represented by matrix 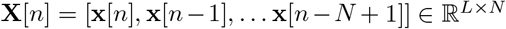. Processing of **X**[*n*] by the first two layers performing spatial and temporal filtering can be described for the m-th branch by a bi-linear product as

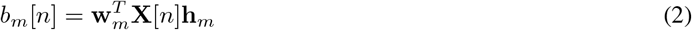

where 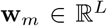 is a vector of spatial weights and 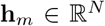 is a vector temporal weights for branch *m*. The non-linearity, *ReLu*(−1), in combination with the low-pass filtering performed by the second convolutional layer (that smooths the rectifier output *r_m_*[*n*]) and extracts the envelopes *e_m_*[*n*] of the rhythmic signals.

We assume that upon training the spatial unmixing coefficients and temporal filter impulse responses implement optimal processing and tune each branch of our architecture to a specific neuronal population with its characteristic geometric and dynamical properties. Under Wiener optimal condition each branch not only gets tuned to a specific population but also tunes itself away from the interfering activity. As detailed in [43], assuming that channel timeseries are zero-mean random processes the underlying neuronal population topographies can be found as

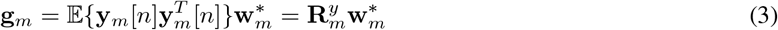

where 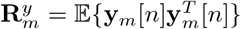 is a *L* × *L* spatial covariance matrix of the temporally filtered data 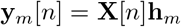, *L* is the number of input channels. Thus, when interpreting individual spatial weights corresponding to each of the *M* branches (or heads) of the architecture shown in Figure 3 one has to take into account the temporal filter weights 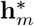 this *m*–th branch is tuned for.

The temporal weights can be interpreted in a similar way taking into account the corresponding spatial filter. Assuming that channel timeseries are zero-mean random processes, *N* is the number of taps in the temporal convolution filter 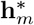, the temporal pattern is given by

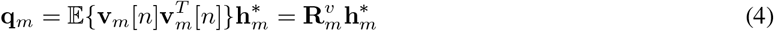

where 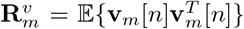 is an *N* × *N* tap covariance matrix of an *N*-samples long chunk of spatially filtered data 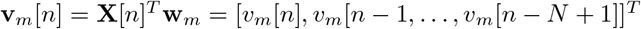.

As shown in [43] if we relax the assumption about the length of the data chunk being equal to the length of the temporal convolution filter we can arrive at Fourier domain representation of dynamics of a neuronal population as pattern *Q_m_*(*f*) derived from the power spectral density (PSD) *P_vm_*(*f*) of the spatially filtered data *v_m_*[*n*] and the Fourier transform *H_m_*(*f*) of the temporal weights vector **h**_m_(*f*) as in 5:

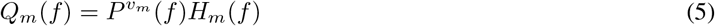

The important distinction that contrasts our weights interpretation approach from the methodology used in the majority of reports utilizing neural networks with separable spatial and temporal filtering operations is that our procedure accounts for the fact that the spatial filter formation is taking place within the context set by the corresponding temporal filter, and vice versa. Also, in [43] the authors for the first time introduced the notion of the frequency domain pattern *Q_m_*(*f*) of neuronal population’s activity. Note that *Q_m_*(*f*) vs. *H_m_*(*f*) has the same difference as spatial pattern vs. spatial filter weights which was brilliantly illustrated earlier in [19].

Using the expressions 3 and 5 we can explore the corresponding spatial and frequency domain patterns of each trained branch (head) of our decoding architecture. If our architecture latched to the data of neuronal origin then the spatial patterns of larger extent should correspond to sources with frequency domain patterns localized to lower frequency ranges and vice versa. Such mutual relationship if observed may reassure that our decoder relies on genuinely neuronal information.

## 4 Results

### 4.1 Microphone effect

To exclude the possibility of data leak associated with electric contacts capacitance change driven by the acoustic speech signal vibes, also known as microphone effect [46], we compared spectral content of the recorded neural data and that of the speech signal in 0-2000 Hz frequency range. Time-frequency diagrams corresponding to a typical 20 seconds long segment of a representative channel of neuronal data and the acoustic signal are shown in Figure 6 for two patients. Visual analysis does not reveal the characteristic banded structure of speech signal (lower row) in the time-frequency profiles of the neuronal data (top row). To make such an objective assessment for all channels of neural data we calculated the correlation coefficient between the temporal profiles of the instantaneous power in each frequency band of neural and acoustic data. We have then used a permutation test to assess statistical significance of the observed correlation coefficients to be non-zero. To this end we split the acoustic data into segments corresponding to word utterances and randomly shuffled *N* = 10000 times the order of such segments to destroy the correspondence between the neuronal and acoustic data in order to compute surrogate correlation coefficient distribution for each (channel, frequency) pair. Then we have computed the asymptotic *p*–values as the fraction of times when the surrogate correlation coefficients appeared to be greater than the correlation coefficients observed in the original non-shuffled data. To correct for multiple comparisons due to running a massive set of tests for all (channel, frequency) pairs we used BH FDR correction procedure [9] and obtained a set of adjusted *p*–values. Those (channel, frequency) pairs whose corresponding adjusted *p*–values fall below 0.05 are highlighted in Figure 5 and do not show any systematic segregation in neither of the two patients.

**Figure 5:**
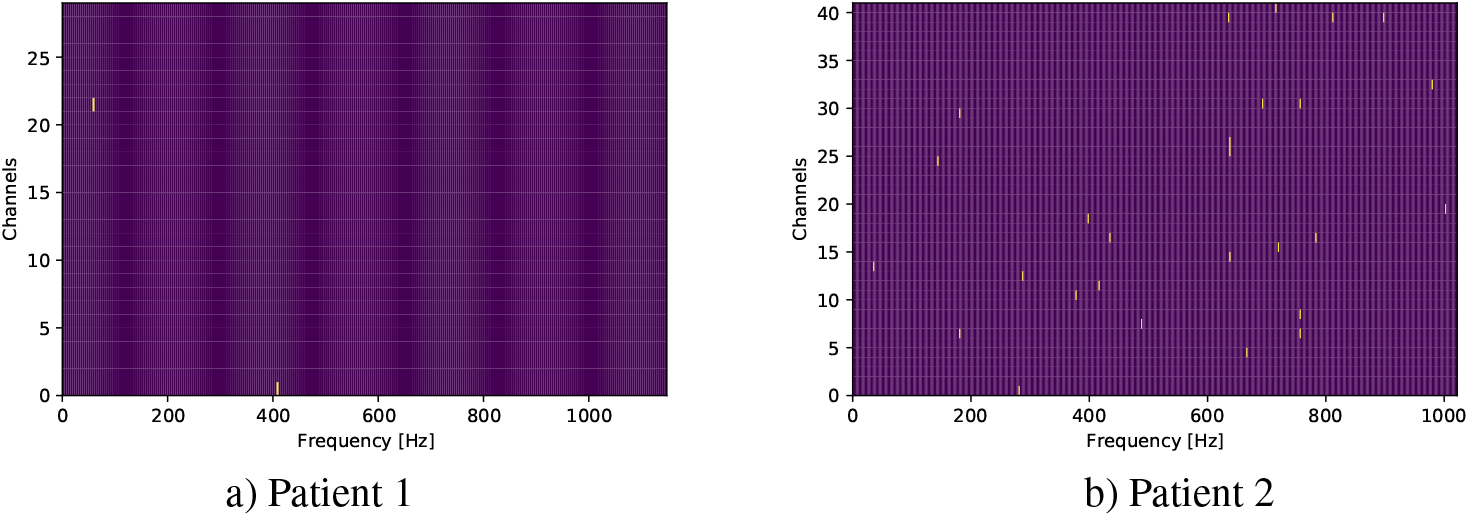
Pairs (channel, frequency) with rejected H0 hypotheses that correlation is zero at *α* = 0.05. See more description in the corresponding section.

**Figure 6:**
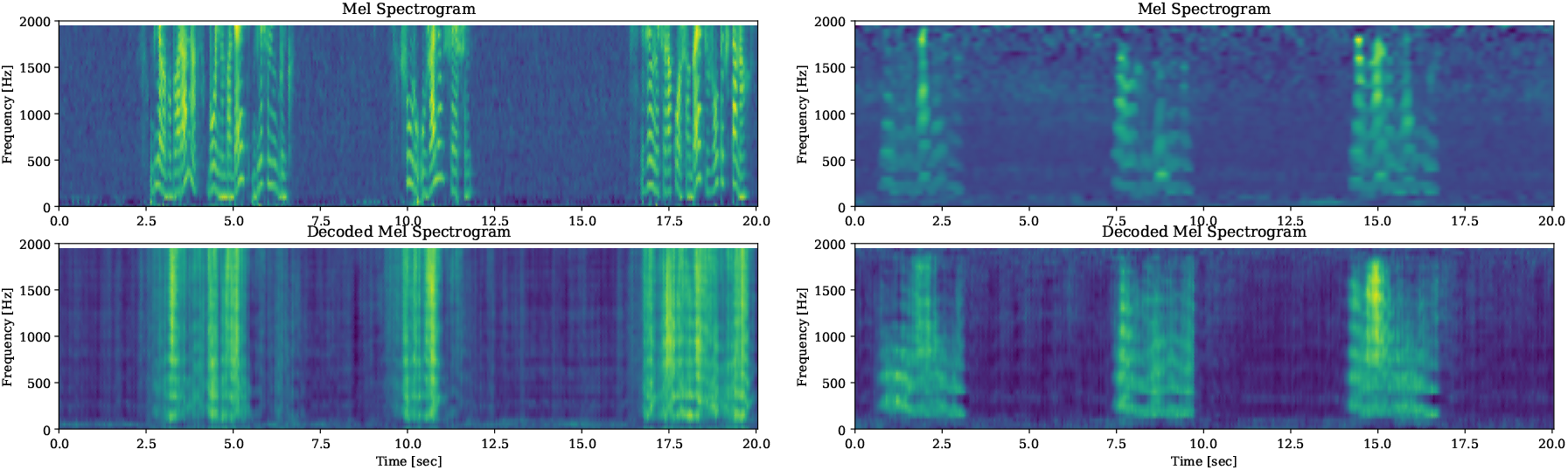
Example of a true and decoded from neural activity log mel-spectrograms.

The above analysis assures that there were no identifiable effects of acoustic information leakage into the data channels carrying neural activity signals.

### 4.2 Decoding internal speech representation

In this study we mainly focused on the contacts confined to a single stereo-EEG shaft in Patient 1 or a single stripe in Patient 2. To select the specific contiguous block of contacts we have computed mutual information between the speech envelope and the envelope of the high-gamma band (60-100 HZ) filtered cortical activity signal for each channel, see Figure 1 c) and d) panels. We have observed a clear delineation in the amount of mutual information between different electrodes. Reassuringly, high MI values closely matched electrodes whose stimulation led to speech arrest in Patient 1 and tongue contraction in Patient 2, see panels a) and b) of Figure 1. Red curves represent the amount of MI computed using the mutually reversed neuronal and audio sequences.

Some remaining values of the MI in the reversed sequence can be explained by the rhythmic structure of the computer instructions to utter that the patients followed.

As evident from Figure 7 our compact architecture using only 6 sEEG channels form a single sEEG shaft achieved about 65% mean correlation over *M* = 40 LMSCs in Patient 1 and almost 60% for Patient 2 with 8 channels from a single ECoG stripe. These accuracy values in decoding internal speech representation are comparable to those reported in [4] where significantly greater count of data channels collected by multiple sEEG shafts was used. An example of the original and decoded 40 LMSCs is shown in Figure 6 for two patients.

**Figure 7:**
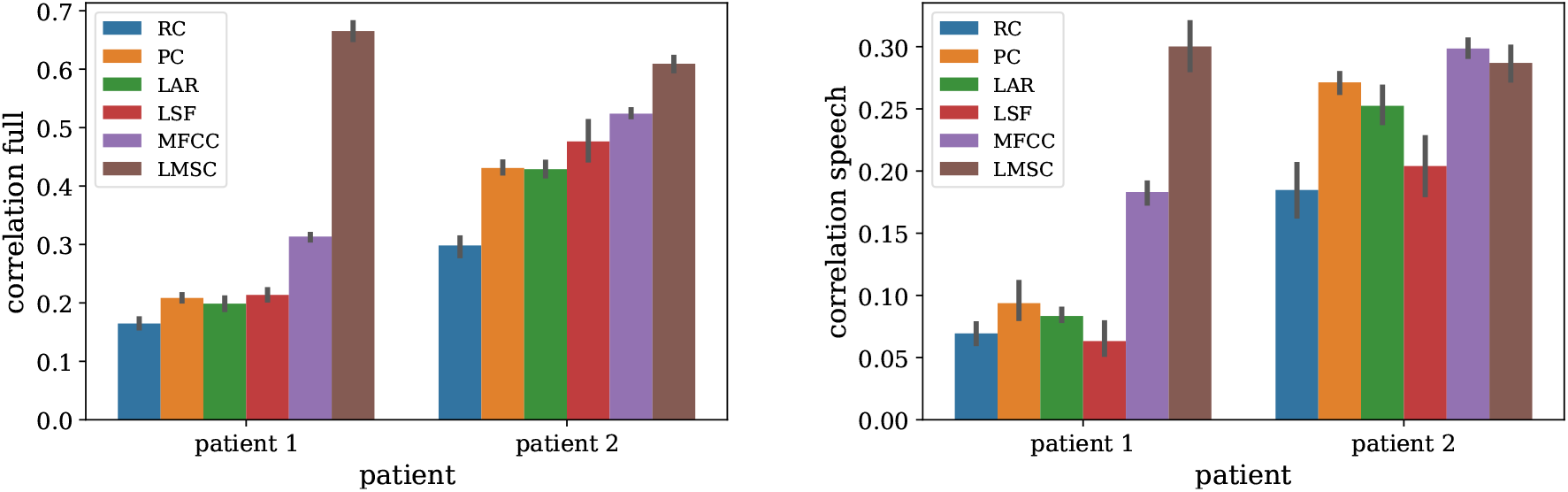
Comparison of the decoding accuracy achieved for different ISRs: PC - autoregressive prediction coefficients, LSF - line spectral frequencies, RC - reflection coefficients, LAR - log-area ratios, LMSCs - log-mel spectrograms, MFCC - mel-frequency cepstral coefficients. The left panel corresponds to the correlation coefficients between the actual and decoded temporal profiles computed over the entire time range of the test data segment. In the right panel the correlation coefficient is computed only over the time intervals where the actual speech was present.

We have also experimented with decoding several other internal speech representations (ISRs) as shown in the left panel of Figure 7. Each color corresponds to a specific ISR method. For both patients we display the ISRs using the same order. Interestingly, in both patients LMSCs appeared to be decoded best, PCs followed and got closely matched by the MFCCs. The reflection coefficients had the worst decoding accuracy. As we will show next, however, this order is not retained when we use words classification accuracy as a criterion. Most likely the mean correlation coefficient between the true and decoded ISRs is determined by their specific statistical properties and the extent to which the fluctuations in their coefficients reflect changes between the silence and speech intervals. To explore this we have computed masked correlation coefficients using only the intervals when the actual speech was present. As expected the mean correlation coefficient dropped significantly and the order in which the different ISRs lined up changed as well, see the right panel of Figure 7. LMSCs still remained among the ISRs with top decoding accuracy in both patients.

Each ISR is a vector and instead of the average values shown in Figure 7 in Figure 8 we present the decoding accuracy values achieved for each of the elements in the three ISRs with the best average decoding accuracy: LMSCs, MFCCs and linear predictive coding PC coefficients. Here we also observe similar tendencies for both patients. For each we show the histograms of correlation coefficients computed over the entire data range (blue) and only over speech intervals (orange).

**Figure 8:**
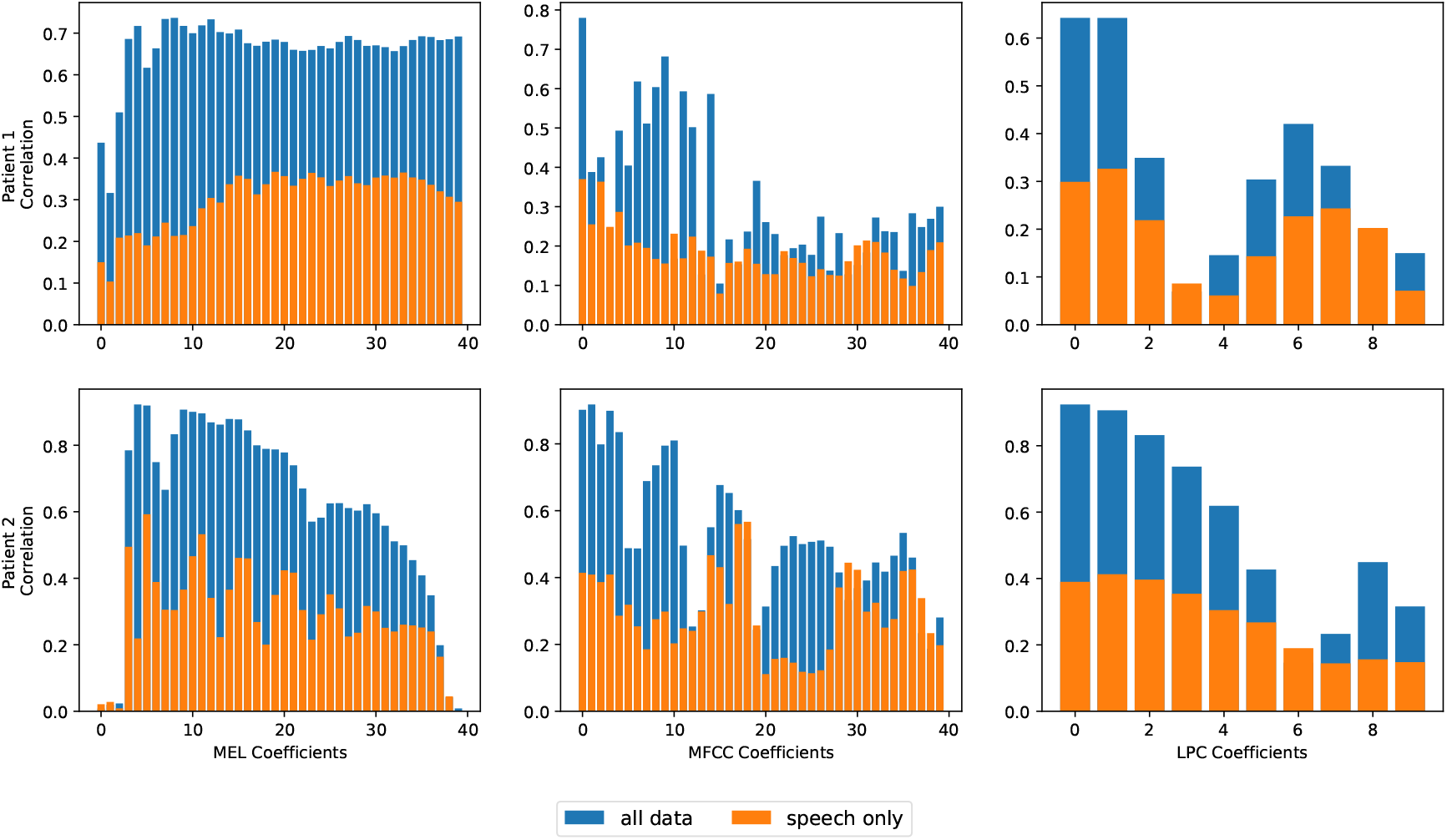
Correlation coefficient of predicted and actual ISR elements for two patients (rows). The two overlayed histogramms correspond to the correlations computed over the entire time range (blue) and only over the speech intervals (orange).

The achieved so far ISR decoding accuracy does not yield intelligible speech when, for example, the recovered LMSC sequence is converted back into the sound. Nevertheless, as we will show next the decoded LMSC profiles and other ISRs support the classification of discrete words sufficiently well.

### 4.3 Words decoding in synchronous mode

We achieved 55% accuracy using only 6 channels of data recorded with a single minimally invasive sEEG electrode in the first patient in classifying 26+1 overtly pronounced words (3.7% chance level). The left panel of Figure 9 shows the corresponding confusion matrix and the individual decoding accuracy values for each word in Patient 1.

**Figure 9:**
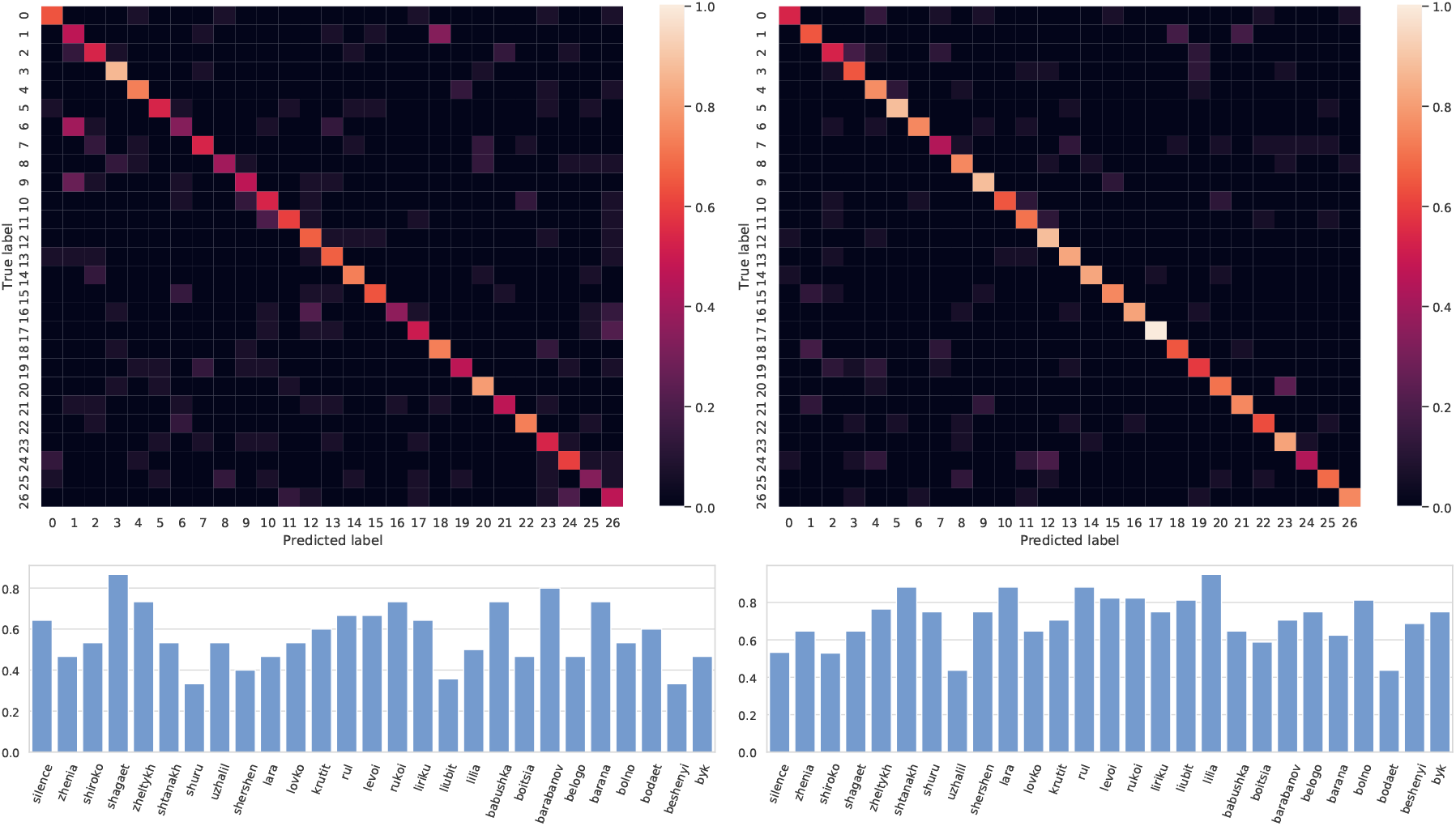
Confusion matrix of classified words for patient 1 and patient 2. Words list: 0. silence, 1. zhenia, 2. shiroko, 3. shagaet, 4. zheltykh, 5. shtanakh, 6. shuru, 7. uzhalil, 8. shershen, 9. lara, 10. lovko, 11. krutit, 12. rul, 13. levoi, 14. rukoi, 15. liriku, 16. liubit, 17. lilia, 18. babushka, 19. boitsia, 20. barabanov, 21. belogo, 22. barana, 23. bolno, 24. bodaet, 25. beshenyi, 26. byk. In the bottom we show the individual word decoding accuracy values, corresponding to the diagonal of the confusion matrix

Spatial characteristics of the first three branches corresponding to the most pivotal neuronal population are shown in the left column of Figure 10.a. We can see that dominantly the activity of these pivotal populations is mapped onto electrodes with indices 9 to 12. This corroborates with the results of an active speech mapping procedure where we found that bipolar electrical stimulation of electrodes indexed 10 and 11 resulted in transient speech arrest as shown in Figure 1 a). Frequency domain patterns presented next to the corresponding spatial patterns illustrate physiological plausibility. First of all the top branch has activity not only in the lower frequency range but also in the traditional gamma band and this branch corresponds to the spatially compact pattern highlighting a single channel with index 12. At the same time, the two branches are characterized by frequency domain patterns concentrated over relatively lower frequency range. Interestingly, and in agreement with [53] these branches have relatively more spread out spatial patterns as compared to that of the first branch.

**Figure 10:**
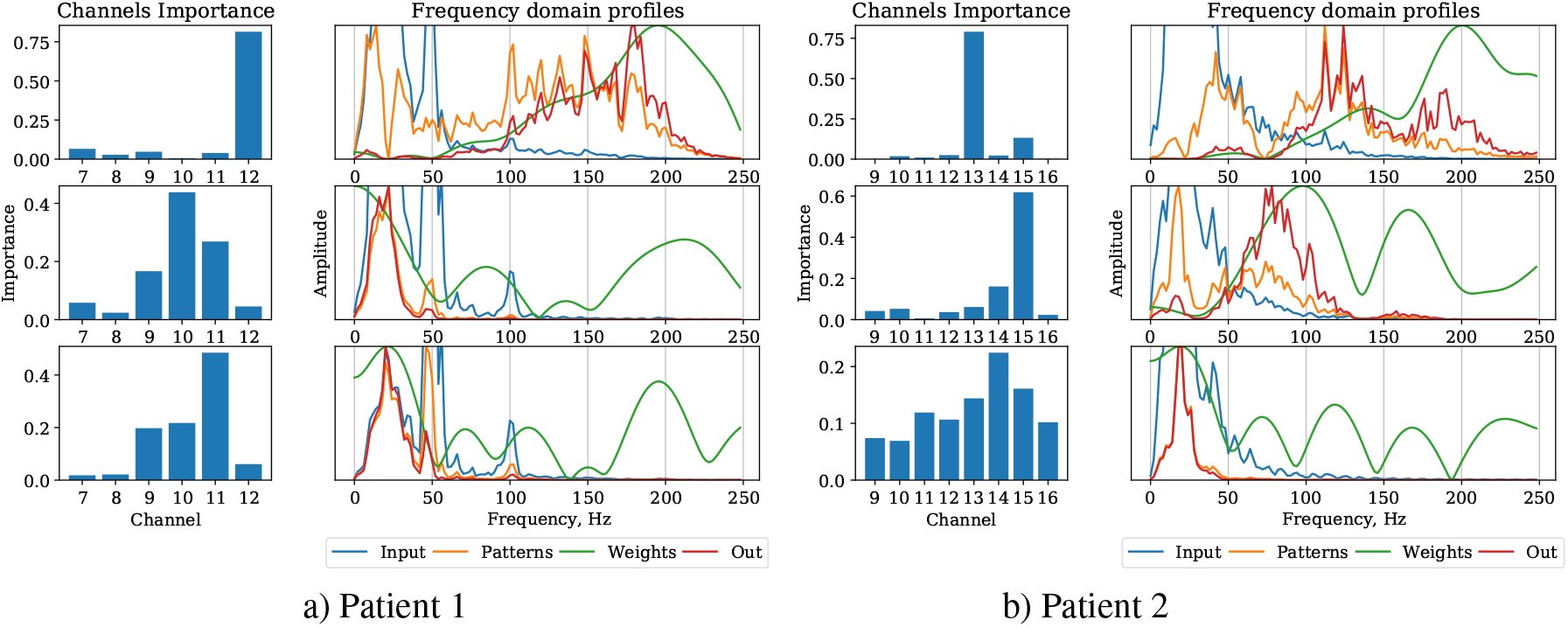
Theoretically justified weights interpretation applied to the most significant branches of the architecture in Figure 3. Orange trace in the top panel shows the power spectral density pattern of the activity of the neuronal population this branch is tuned to. The left panel shows the spatial pattern of this population. We can conclude that this source dominantly projects onto the 12th contact located at the lateral part of the sEEG electrode (shaft), see Figure 1 a). Similar picture is observed for the second patient present in the two right columns. Here we also observe a striking trend of spatially more compact populations being characterized by the activity in the higher frequency bands.

Similar analysis is shown for Patient 2 in the right panel of Figure 9 and Figure 10.b. In this patient implanted with ECoG grids we have achieved on average 70% of words decoding accuracy. We can also observe a striking trend where spatially more compact populations are characterized by the activity in the higher frequency bands.

#### 4.3.1 Weights interpretation

The advantage of the DNN based approach is that it does not require manual feature engineering, however, these methods are typically over-parameterized and exhibit greedy behaviour. Such a greediness in the neurophysiological context may result in the network latching on signals of non-neuronal origin. This problem can be monitored in compact, domain knowledge driven architectures equipped with a proper weights interpretation approach. To this end we have applied the recently developed approach detailed in [43]. Our goal here is to explore the mutual relation between the spatial and frequency domain patterns each branch of our compact DNN architecture got tuned to during the training process. This analysis will also help us to exclude the fact that our network exploits muscular activity associated with the speech production process. The principles behind this analysis have been briefly outlined in section 3.6.

The result of applying our weights interpretation procedure to each of the three branches of our compact DNN is shown in Figure 10. We illustrate both spatial (left column) and frequency domain (right column) patterns of the neuronal populations for each of the three most significant branches of our network. The frequency domain plot also contains the curves corresponding to the power spectral density (PSD) of the input timeseries obtained by the spatial filtering of the multichannel data at the input of the network and the PSD of the branch’s output timeseries.

From the top row of patterns corresponding to the first branch of the decision rule for Patient 1 we can see that the PSD occupies high 100-200 Hz frequency range and the corresponding spatial pattern is confined to only a single channel with index 12. At the same time the second branch with a much more spread out spatial pattern occupying channels 9-11 is characterized by the PSD confined to the lower 10-40 Hz frequency range. The reciprocal space-frequency relation that hallmarks neuronal activity and distinguishes it from the electro-muscular artifacts is also very well pronounced in the second patient. Moving downwards we observe the gradual growth of the spatial spread with the PSD frequency range migrating from the higher to lower frequency range.

Combined together with domain knowledge [11, 12, 10, 53] highlighting reciprocal space-time relationship in the observed cortical activity patterns and phenomenological observations [15] on the properties of the electromuscular activity and its representation on the cortex the observed combinations of the spatial and PSD patterns allow us to make a conclusion regarding the neuronal origin of the data our decoder latched on during the training process. The analysis for microphone effect reported in section 4.1 also excludes the possibility that the decoding is done based on the acoustic signal leaking into neuronal data channels.

In this patient we have witnessed certain discrepancy between the stimulation based mapping and the electrode indexes that resulted from our weights interpretation procedure where electrode 12 was highlighted, yet no speech production problems were registered when stimulating contacts 11-12, see Figure 1 a,c. This could have resulted from the very sparing stimulation settings used in this patient - our stimulation current never exceeded 3 mV which is below the traditional average current magnitude typically used for speech mapping [14].

For Patient 2, weights interpretation of the three most important branches of our network show the primary involvement of electrodes 13, 14 and 15 into the decoding process which is partially congruent with stimulation based speech mapping, see Figure 1.b,d, where we found that the stimulation applied between 15-16th electrodes yielded reliable tongue retraction (back from the requested tongue protrusion state). In this patient we also observe a very pronounced reciprocity in space-frequency patterns. As shown in the right panel of Figure 10 moving from the top most spatially compact pattern to the spatially spread out profile of the third branch shown in the bottom we observe the expected migration of the frequency domain pattern from the high to the lower frequency range

Approximate MNI coordinates of electrodes found to be pivotal for decoding in both patients are given in Table 1. In Patient 1 stereo-EEG electrodes with the majority of contacts located deep in the sulci of the left operculum. The location corresponds to Brocas region whose activity is traditionally registered in a broad range of language related tasks. For patient 2 the pivotal electrodes cover the inferior portion of the precentral gyrus. Our stimulation results in Patient 2 do not quite match the anatomical location and better correspond to the ventral precentral gyrus, the structure located inferior to the precentral gyrus and known to house the tongue motor area. This could be due to atypical organization of the cortex in this patient. Locations of electrodes in both patients are remote with respect to the belt area (MNI: −58, −28, 13) whose gamma-band activity was shown to reliably track the perceived speech envelope (Kubanek et al., 2013). These functional and anatomical arguments together with the causal approach to the ISR decoding reduce the chance that our decoder operation is based on the subjects own speech perception.

**Table 1:** MNI coordinates of pivotal electrodes

| Patient 1, stereo-EEG electrode |  | Patient 2, ECoG strip |  |
| --- | --- | --- | --- |
| Electrode index | MNI Coordinates | Electrode index | MNI Coordinates |
| 9 | -40,19,4 | 13 | -65,-24,43 |
| 10 | -45,20,4 | 14 | -66,-18,39 |
| 11 | -50,20,4 | 15 | -67,-13,34 |
| 12 | -53,21,4 |  |  |

In the above we have analyzed spatial and frequency domain patterns of the neuronal populations that were found to be pivotal to the ISR decoding task and forming the internal representations to be subsequently used as an input to our words classification network. For the weights interpretation to make sense we also need to demonstrate the dependence of the final word classification accuracy on the individual ISR decoding fidelity. To this end we have performed additional experiments. We rerun the training and terminated it at different points to yield various ISR decoding accuracy and then subsequently trained word classification network. In Figure 12 we show the observed dependence of the words classification accuracy on the correlation coefficient between the actual and the decoded ISRs (LMSC, MFCC and LPC). Indeed for each ISR we witness the direct relation between its decoding fidelity as measured by the correlation coefficient and the corresponding discrete words classification accuracy.

At the same time, although the ISRs differed in the quality of their decoding, see Figure 7 these differences were not carried over to the final words classification accuracy, see Figure 11. A possible explanation here could be that some ISRs in addition to the information regarding the sequence of the articulatory tract configurations (corresponding to a specific sequence of phonemes and invariant to the pitch, timbre, loudness, etc.,) contain the information about purely acoustic features of the utterance such as fundamental frequency, voice timbre, local volume, etc., which could be easier to decode than the articulatory tract parameters critical for the words classification task. The subsequent words classification largely requires only the first type of information and therefore may yield comparable performance for the different ISRs as long as all of them contain this essential information. In Figure 12 we can observe very characteristic shapes in the depicted curves. For ISR = LMSC, the increase of the ISR decoding accuracy up to 0.4 in both patients does not yield any tangible improvement in the words classification accuracy. This could be due to the fact that the first 40% of correlation between the recovered and actual LMSC profiles is achieved by reproducing purely acoustic features present in the LMSC (or even simply decoding on-off states) not directly related to the specific uttered word. Only advancing LMSC decoding accuracy further do we get the expected increase in words classification. This is likely to indicate that this further increase was achieved due to the front-end network starting to learn content dependent information present in the LMSC. At the same time, it is known that both MFCC and LPC primarily encode the articulatory tract dynamics and are largely invariant to the speech acoustic features. For these ISRs we observe a somewhat earlier start of the ascent in the depicted curves indicating that their decoding accuracy more rapidly gets translated into the improvement of words classification.

**Figure 11:**
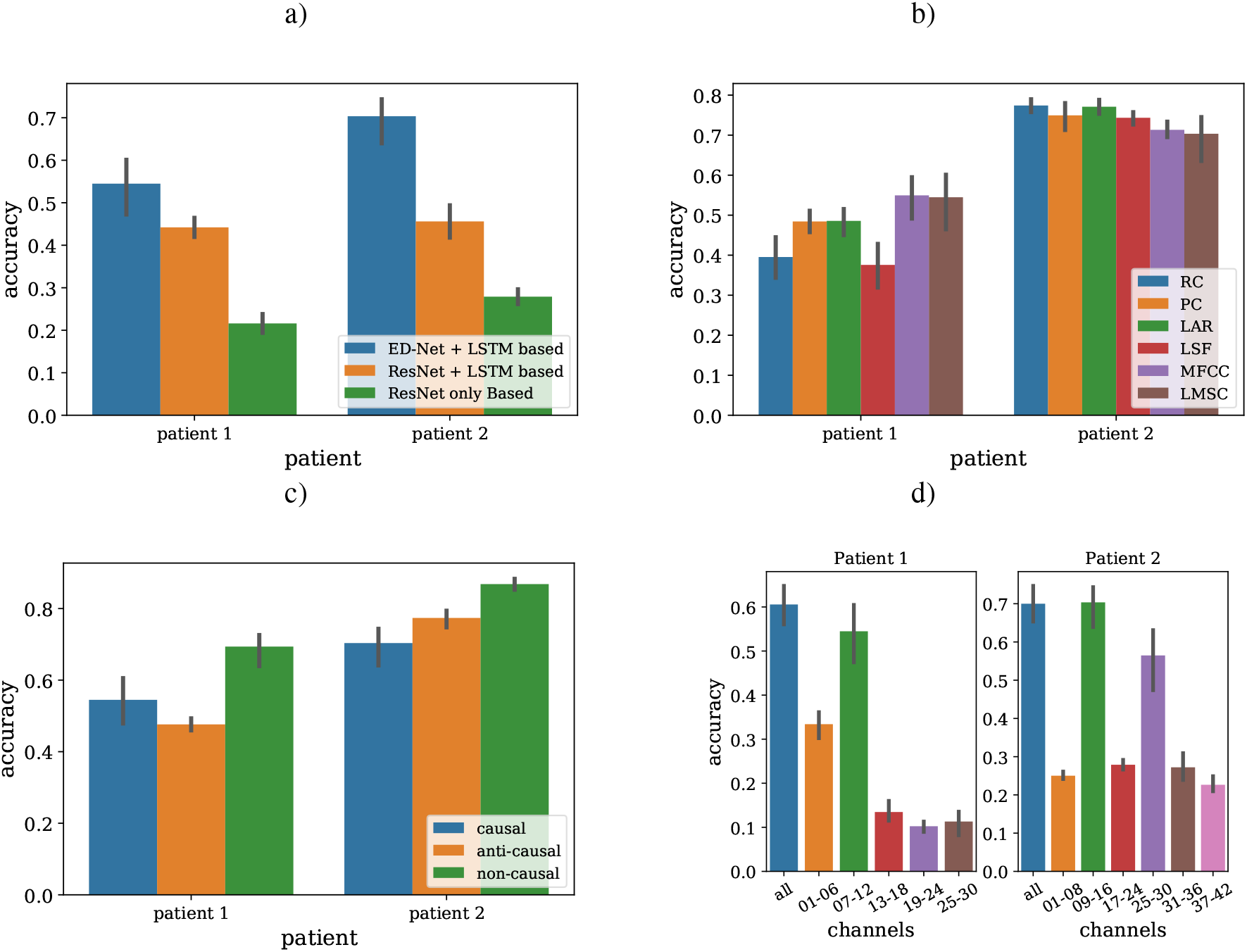
Comparative analysis. a) Comparison of different neural network models b) Comparison of different possible intermediate sound representation, PC − autoregressive prediction coefficients, LSF - Line Spectral Frequencies, RC - reflection coefficients, LAR - log-area ratios, LMSC - log-mel spectrogram coefficients, MFCC - mel-frequency cepstral coefficients c) Comparison of different possible lag d) Comparison of decoding quality for different subset of channels.

**Figure 12:**
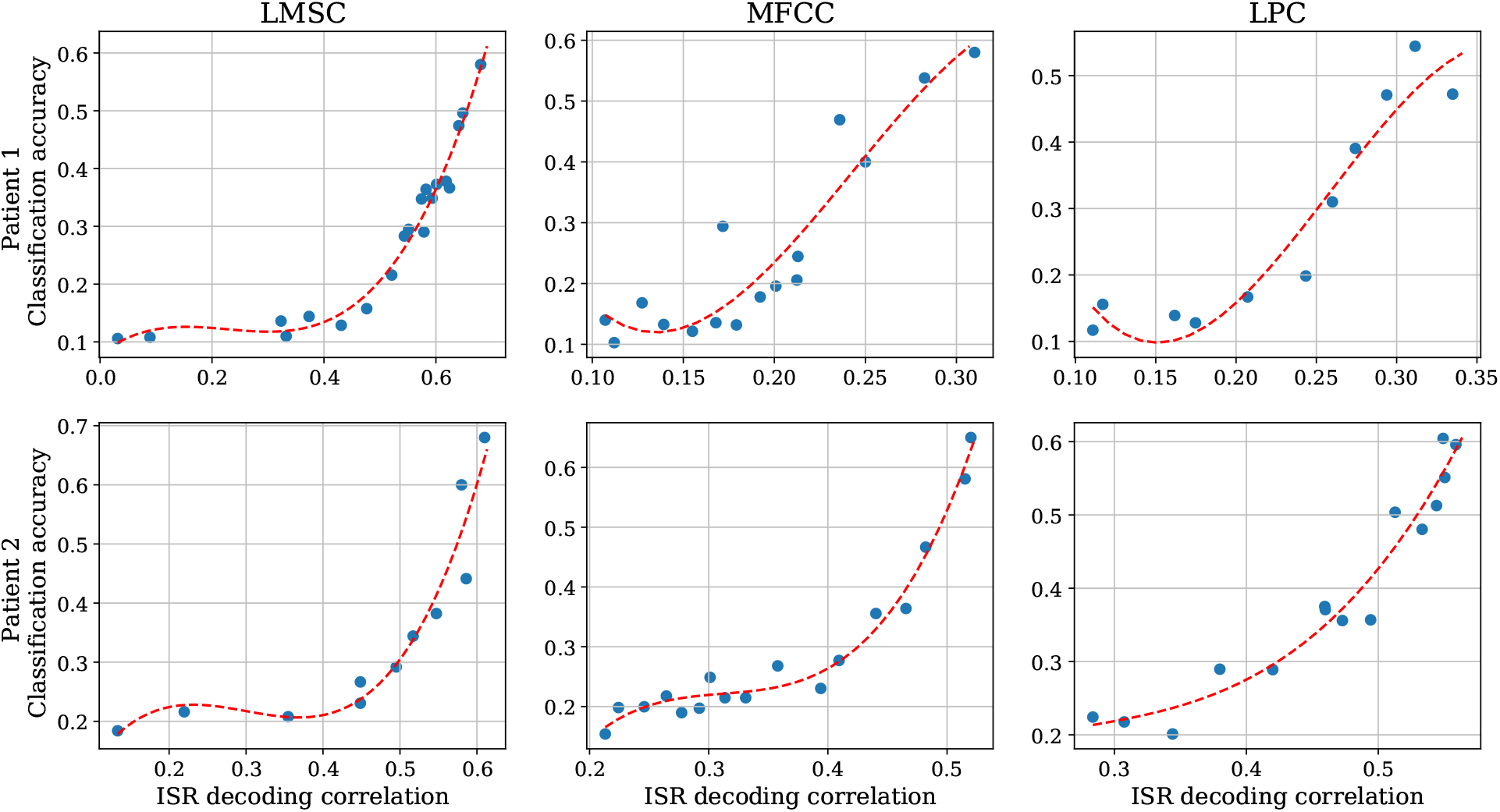
Dependence of the final word classification accuracy on the ISR decoding quality. Red line is the third order trend fitted to the data points to facilitate visual perception.

### 4.4 Comparative analysis

In this work we employed the compact architecture, see Figure 3 that comprises multiple branches of envelope detectors (ED) of spatially filtered data whose output is fed into the LSTM layer followed by a fully connected network. This architecture uses factorized spatial and temporal filters that get adapted during training and allows for interpretation of the filter weights into the spatial and spectral patterns as demonstrated in Figure 10. These patterns can then be used to infer location and dynamical properties of the underlying neuronal populations.

Here we compared this network to several other architectures. We found that out of several neural networks only Resent-18 offers a comparable, although significantly worse, performance when used instead of the ED block in our architecture, see Figure 3. The LSTM layer also appears to be very useful in capturing the dynamics of features extracted either with ED or ResNet blocks, see Figure 11.a. We hypothesize that this situation may be caused by the adequate balance in the number of parameters to be tuned for the ED-based network and the amount of data available for training as compared to several other more sophisticated architectures.

Words decoding accuracy results reported in Figure 9 correspond to the case when 40 LMSCs were used to train the front-end ISR decoder network, see Figure 3. We have also experimented with several other ISRs as described in section 3.3 and presented the results in Figure 11.b. We can see that for the first patient log-mel spectrum coefficients (LMSC) target results into the highest words decoding accuracy. Interestingly, in contrast to the actual ISR decoding task displayed in Figure 7 the difference in the words decoding accuracy between various ISRs seems to be significantly less articulated than the differences in the quality of decoding of each of such representations. Nevertheless for both patients we observe the similar pattern with PC and LSF yielding relatively worse words decoding accuracy than the other ISRs. In this analysis LPC reflection coefficients (RC) yield better decoding accuracy as compared to the prediction coefficients. This observation matches the properties of the RC coefficients as informationally equivalent but a more stable version of the original PC.

The reported ISR and words decoding accuracy results are presented for the causal processing mode, i.e. when the data window strictly precedes the time-point the prediction is made for. We have also experimented with anti-causal (the window is strictly in the future w.r.t. to the predicted time-point) and non-causal (when the data window covers pre- and post-intervals around the point in question). These results are plotted in Figure 11.c. In both patients we see the best performance when the data-window is allowed to be located both in the future and in the past w.r.t. the point to be predicted. This result is expected since in the non-causal setting the algorithm can use information about the cortical activity that occurs in response to the uttered word.

In this work we mainly focused on decoding from a small number of contacts confined either to a single stereo-EEG shaft or an ECoG stripe. In both cases the electrodes can be implanted without a full-blown craniotomy via a drill hole in the skull. We have chosen the particular subset of contacts using the mutual information (MI) metric, see Figure 1 which closely matched stimulation-based mapping results. Both of our patients were implanted with several sEEG shafts or ECoG stripes, see Figure 1. In Figure 11.d we show the results of a similar analysis but using other subsets of electrodes located on the other shafts or stripes. Noteworthy is that MI based selection yielded significantly better performance as compared to the other spatially segregated electrode groups.

### 4.5 Asynchronous decoding of words

Traditionally, BCI can be used in two different settings: synchronous and asynchronous. In the synchronous setting a command is to be issued within a specific time window. Usually, a synchronous BCI user is prompted at the start of such a time window and has to produce a command (alter his or her brain state) within a specified time frame. Therefore, the decoding algorithm is aware of the specific segment of data to process in order to extract the information about the command. In the asynchronous mode the BCI needs to not only decipher the command but also determine the fact that the command is actually being issued. The delineation between synchronous and asynchronous modes is most clearly pronounced in BCIs with discrete commands implying the use of a categorical decoder.

In BCIs that decode a continuous variable, e.g. hand kinematics, such delineation between synchronous and asynchronous modes is less clear. The first part of our BCI implements a continuous decoder of the internal speech representation (ISR) features. Should this decoding appear of sufficient accuracy it could have been simply used as an input to a voice synthesis engine. Such scenario has already been implemented in several reports [4, 3] but these solutions use a large number of electrodes which may explain better quality of ISR decoding. In our setting we aimed at building a decoder operating with a small number of ecologically implanted electrodes and decided to focus on decoding individual words. We first used the continuously decoded ISRs to classify 26 discrete words and one silence state in the synchronous manner. To implement this we cut the decoded ISR timeseries around each word’s utterance and use them as data samples for our classification engine.

To gain insight into the ability of our BCI to operate in a fully asynchronous mode we performed the additional analysis as described in section 3.3.2. Figure 13 b) illustrates the performance of our BCI operating in a fully asynchronous mode when the decoder is running over the succession of overlapping time windows of continuously decoded ISRs and the decision about the specific word being uttered is made for each of such windows, see Figure 2. To quantify the performance of our asynchronous speech decoder we used precision-recall curves as detailed in section 3.5 and Figure 13 a).

**Figure 13:**
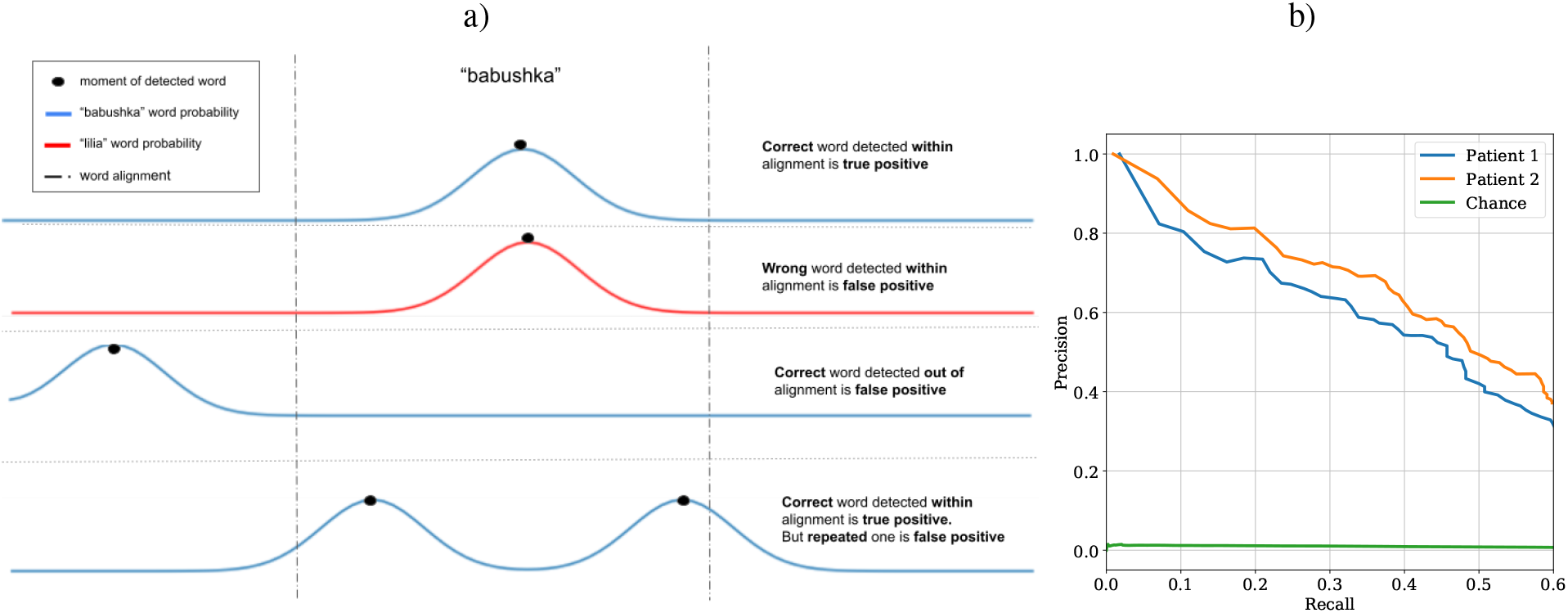
a) For each *i*–th word we compute smoothed probability profiles 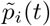 for each time instance *t*. The decision is then made about a word being pronounced only at time points corresponding to the local maximums of 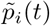 that cross the threshold *θ*. In case the chosen *i*–th word matches the one that is currently being uttered we mark this event as true positive (TP). If after such a detection 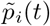 remains above the threshold and exhibits another local maximum which exceeds the values of all other smoothed probability profiles the *i* – th word is “uttered” again, but this event is marked as false positive (FP) even if *t* belongs to the time range corresponding to the actual *i* – th word. b) PR curves for asynchronous words decoding task. Note that definition of precision and recall is slightly different from conventional binary classification PR curves (see equation 5). We also show a chance level PR curve.

Although the observed performance significantly exceeds the chance level, it is not yet sufficient for building a full blown asynchronous speech interface operating using a small number of minimally invasive electrodes. In our view and based on the experience with motor interfaces, specific protocols to train the patient including those with immediate feedback to the user [6] are likely to significantly improve the decoding accuracy in such systems which will boost the overall feasibility of minimally invasive speech prosthetic solutions.

## 5 Conclusion

We have explored the possibility of building a practically feasible speech prosthesis solution operating on the basis of neural activity recorded with a small set of minimally invasive electrodes. Implantation of such electrode systems does not require a full craniotomy and combined with algorithmic solutions equipped with a joint human-machine training protocol may form a basis for the future minimally invasive speech prosthesis.

There exist several reports exploiting intracortical activity recorded with Utah array like systems for speech prosthesis purposes [56, 54, 28]. These recordings give access to the activity of individual neurons but remain potentially harmful to the cortical tissue. In contrast, stentrodes [41], electrodes located inside blood vessels and implanted using stent technology, offer a potentially plausible solution for obtaining high quality brain activity signals without any kind of craniotomy. These electrodes, however, unlike the intracortical arrays, register the superposition of neuronal activity stemming from a large number of neuronal populations. Also, unlike the ECoG grids used in the majority of speech prosthesis research these stent electrodes are confined to a relatively small volume. The signals measured in our setting with a small number of spatially confined sEEG and ECoG contacts can be considered as a proxy of the data collected by the stentrodes and the signal processing approaches developed here could be potentially applied to stentrode data in order to to pave the road towards craniotomy-free speech BCI solutions.

We build our decoder using a two-step procedure. First we construct an interpretable architecture to decode the continuous internal speech representation (ISR) profiles from the neural activity and fix the weights of this compact neural network. In this case the particular ISR (LMSC, MFCC, LPC coefficients) is merely a target to train this front-end network. Then, when applying this network to neural activity data we take its hidden state before the last fully connected layer and use it as an input to the discrete classifier to distinguish between neural activity patterns corresponding to 26 words and one silence state. This approach resembles [34]. However, based on our experiments we found that replacing concurrent training of two classifiers with such a two step process in our setting improves the achieved decoding accuracy.

We have also paid particular attention to interpreting the obtained decision rule. Our main concern here was to exclude the possibility of using non-neural activity patterns in the overt speech decoding setting. To do so we exploited the concept of spatial and frequency domain patterns that pertain to the neuronal populations that each of the branches of our front-end network got tuned to. Several reports [15, 29, 7] explored the spatial and frequency domain patterns that manifest muscular activity in the subdural space. These are typically hallmarked with high-frequency spectra and large spatial extent which is the opposite to neural activity where we expect higher frequency activity to be more spatially confined as compared to the signals in the lower frequency bands. We applied the methodology described in [43] to recover spatial and frequency patterns of the underlying pivotal activity and found that they well adhered to the described properties of neural activity. We also did not find any evidence of microphone effect [45] in our data.

The accuracy we obtained in the synchronous mode appears sufficient to make the system usable in a real-life scenario when each word is “uttered” within a specific time slot, starting, for example, with a beep prompt. The extent to which the observed accuracy is transferred to a patient who lacks the ability to speak greatly depends on the specific medical case. Although we explored various arrangements of the data time window around the decision point our main results correspond to the decoder operating causally, i.e. utilizing neural activity strictly from the past which is expected not to depend on the perceived speech. This ensures that the observed accuracy will transfer to real patients with speech function deficits given the appropriate training tools for patients are developed.

Asynchronous BCI setting is clearly a more natural one for speech prosthesis operation. We experimented with our decoder in this scenario and observed a reasonable performance which however, needs to be improved before it can be used in practice. We recall 40% moments when one of the 26 words is uttered it was also detected by our approach and in 60% of cases we correctly guessed this word out of 26 possible alternatives.

The use of a language model is known to improve speech decoding accuracy [51] and can also be added to improve the accuracy of the final consumer solution. However, our goal here was to assess to which extent the neural activity along can be informative with regard to individual words classification and therefore we have deliberately refrained from using any language model in this study.

Overall our study showcases the possibility of building speech prosthesis with a small number of electrodes and based on a compact feature engineering free decoder derived from several tens of minutes worth of training data. To be translated into clinical practice this solution needs to be augmented with patient training procedures and a methodology to non-invasively determine implantation sites that would yield the best speech decoding accuracy.

## Acknowledgment

This work is supported by the Center for Bioelectric Interfaces NRU HSE, RF Government grant, AG. No. 075-15-2021-624

## References

[1] Sarah N Abdulkader, Ayman Atia, and Mostafa-Sami M Mostafa. Brain computer interfacing: Applications and challenges. Egyptian Informatics Journal, 16(2):213–230, 2015.

[2] Abidemi B Ajiboye and Robert F Kirsch. Invasive brain–computer interfaces for functional restoration. In Neuromodulation, pages 379–391. Elsevier, 2018.

[3] Hassan Akbari, Bahar Khalighinejad, Jose L Herrero, Ashesh D Mehta, and Nima Mesgarani. Towards reconstructing intelligible speech from the human auditory cortex. Scientific reports, 9(1):1–12, 2019.

[4] Miguel Angrick, Christian Herff, Emily Mugler, Matthew C Tate, Marc W Slutzky, Dean J Krusienski, and Tanja Schultz. Speech synthesis from ecog using densely connected 3d convolutional neural networks. Journal of neural engineering, 16(3):036019, 2019.

[5] Miguel Angrick, Maarten Ottenhoff, Lorenz Diener, Darius Ivucic, Gabriel Ivucic, Sofoklis Goulis, Jeremy Saal, Albert J Colon, Louis Wagner, Dean J Krusienski, et al. Real-time synthesis of imagined speech processes from minimally invasive recordings of neural activity. bioRxiv, 2020.

[6] Miguel Angrick, Maarten Ottenhoff, Lorenz Diener, Darius Ivucic, Gabriel Ivucic, Sophocles Goulis, Albert J Colon, Louis Wagner, Dean J Krusienski, Pieter L Kubben, et al. Towards closed-loop speech synthesis from stereotactic eeg: A unit selection approach. In ICASSP 2022-2022 IEEE International Conference on Acoustics, Speech and Signal Processing (ICASSP), pages 1296–1300. IEEE, 2022.

[7] Tonio Ball, Markus Kern, Isabella Mutschler, Ad Aertsen, and Andreas Schulze-Bonhage. Signal quality of simultaneously recorded invasive and non-invasive eeg. Neuroimage, 46(3):708–716, 2009.

[8] Kalaba Bellman. Bellman r., kalaba r. On adaptive control processes, IRE Trans. Autom. Control, 4(2):1–9, 1959.

[9] Yoav Benjamini and Yosef Hochberg. Controlling the false discovery rate: a practical and powerful approach to multiple testing. Journal of the Royal statistical society: series B (Methodological), 57(1):289–300, 1995.

[10] Peter Brunner, Anthony L Ritaccio, Timothy M Lynch, Joseph F Emrich, J Adam Wilson, Justin C Williams, Erik J Aarnoutse, Nick F Ramsey, Eric C Leuthardt, Horst Bischof, et al. A practical procedure for real-time functional mapping of eloquent cortex using electrocorticographic signals in humans. Epilepsy & Behavior, 15(3):278–286, 2009.

[11] Gyorgy Buzsaki. Rhythms of the Brain. Oxford University Press, 2006.

[12] György Buzsáki, Costas A Anastassiou, and Christof Koch. The origin of extracellular fields and currentseeg, ecog, lfp and spikes. Nature reviews neuroscience, 13(6):407–420, 2012.

[13] Ujwal Chaudhary, Niels Birbaumer, and Ander Ramos-Murguialday. Brain–computer interfaces for communication and rehabilitation. Nature Reviews Neurology, 12(9):513, 2016.

[14] Jacquelyn A. Corley, Pouya Nazari, Vincent J. Rossi, Nora C. Kim, Louis F. Fogg, Thomas J. Hoeppner, Travis R. Stoub, and Richard W. Byrne. Cortical stimulation parameters for functional mapping. Seizure, 45:36–41, 2017.

[15] Andrey Eliseyev and Tatiana Aksenova. Stable and artifact-resistant decoding of 3d hand trajectories from ecog signals using the generalized additive model. Journal of neural engineering, 11(6):066005, 2014.

[16] Michael J Fagan, Stephen R Ell, James M Gilbert, E Sarrazin, and Peter M Chapman. Development of a (silent) speech recognition system for patients following laryngectomy. Medical engineering & physics, 30(4):419–425, 2008.

[17] Joris Guérin, Stephane Thiery, Eric Nyiri, Olivier Gibaru, and Byron Boots. Combining pretrained cnn feature extractors to enhance clustering of complex natural images. Neurocomputing, 423:551–571, 2021.

[18] Nicholas G Hatsopoulos and John P Donoghue. The science of neural interface systems. Annual review of neuroscience, 32:249–266, 2009.

[19] Stefan Haufe, Frank Meinecke, Kai Görgen, Sven Dähne, John-Dylan Haynes, Benjamin Blankertz, and Felix Bießmann. On the interpretation of weight vectors of linear models in multivariate neuroimaging. Neuroimage, 87:96–110, 2014.

[20] Christian Herff, Lorenz Diener, Miguel Angrick, Emily Mugler, Matthew C Tate, Matthew A Goldrick, Dean J Krusienski, Marc W Slutzky, and Tanja Schultz. Generating natural, intelligible speech from brain activity in motor, premotor, and inferior frontal cortices. Frontiers in neuroscience, 13:1267, 2019.

[21] Christian Herff, Dean J Krusienski, and Pieter Kubben. The potential of stereotactic-eeg for brain-computer interfaces: current progress and future directions. Frontiers in neuroscience, 14:123, 2020.

[22] Sepp Hochreiter and Jürgen Schmidhuber. Long short-term memory. Neural computation, 9(8):1735–1780, 1997.

[23] Mark L Homer, Arto V Nurmikko, John P Donoghue, and Leigh R Hochberg. Sensors and decoding for intracortical brain computer interfaces. Annual review of biomedical engineering, 15:383–405, 2013.

[24] Gao Huang, Zhuang Liu, Laurens Van Der Maaten, and Kilian Q Weinberger. Densely connected convolutional networks. In Proceedings of the IEEE conference on computer vision and pattern recognition, pages 4700–4708, 2017.

[25] Xuedong Huang, Alex Acero, Hsiao-Wuen Hon, and Raj Reddy. Spoken Language Processing: A Guide to Theory, Algorithm, and System Development. Prentice Hall PTR, USA, 1st edition, 2001.

[26] Prasanna Jayakar, Jean Gotman, A. Simon Harvey, André Palmini, Laura Tassi, Donald Schomer, Francois Dubeau, Fabrice Bartolomei, Alice Yu, Pavel Krek, Demetrios Velis, and Philippe Kahane. Diagnostic utility of invasive eeg for epilepsy surgery: Indications, modalities, and techniques. Epilepsia, 57(11):1735–1747, 2016.

[27] Rachel Kaye, Christopher G Tang, and Catherine F Sinclair. The electrolarynx: voice restoration after total laryngectomy. Medical Devices (Auckland, NZ), 10:133, 2017.

[28] Philip Kennedy, A Ganesh, and AJ Cervantes. Slow firing single units are essential for optimal decoding of silent speech. 2022.

[29] Christopher K Kovach, Naotsugu Tsuchiya, Hiroto Kawasaki, Hiroyuki Oya, Mathew A Howard III, and Ralph Adolphs. Manifestation of ocular-muscle emg contamination in human intracranial recordings. Neuroimage, 54(1):213–233, 2011.

[30] Jan Kubanek, Peter Brunner, Aysegul Gunduz, David Poeppel, and Gerwin Schalk. The tracking of speech envelope in the human cortex. PloS one, 8(1):e53398, 2013.

[31] Mikhail A Lebedev and Miguel AL Nicolelis. Brain-machine interfaces: From basic science to neuroprostheses and neurorehabilitation. Physiological reviews, 97(2):767–837, 2017.

[32] Sergio Machado, Fernanda Araújo, Flávia Paes, Bruna Velasques, Mario Cunha, Henning Budde, Luis F Basile, Renato Anghinah, Oscar Arias-Carrión, Mauricio Cagy, et al. Eeg-based brain-computer interfaces: an overview of basic concepts and clinical applications in neurorehabilitation. Reviews in the Neurosciences, 21(6):451–468, 2010.

[33] Joseph N Mak and Jonathan R Wolpaw. Clinical applications of brain-computer interfaces: current state and future prospects. IEEE reviews in biomedical engineering, 2:187–199, 2009.

[34] Joseph G Makin, David A Moses, and Edward F Chang. Machine translation of cortical activity to text with an encoder–decoder framework. Nature Neuroscience, 23(4):575–582, 2020.

[35] L. Marple. A new autoregressive spectrum analysis algorithm. IEEE Transactions on Acoustics, Speech, and Signal Processing, 28:441–454, 1980.

[36] Brian McFee, Colin Raffel, Dawen Liang, Daniel P Ellis, Matt McVicar, Eric Battenberg, and Oriol Nieto. librosa: Audio and music signal analysis in python. In Proceedings of the 14th python in science conference, volume 8, pages 18–25. Citeseer, 2015.

[37] David A Moses, Sean L Metzger, Jessie R Liu, Gopala K Anumanchipalli, Joseph G Makin, Pengfei F Sun, Josh Chartier, Maximilian E Dougherty, Patricia M Liu, Gary M Abrams, et al. Neuroprosthesis for decoding speech in a paralyzed person with anarthria. New England Journal of Medicine, 385(3):217–227, 2021.

[38] Emily M Mugler, James L Patton, Robert D Flint, Zachary A Wright, Stephan U Schuele, Joshua Rosenow, Jerry J Shih, Dean J Krusienski, and Marc W Slutzky. Direct classification of all american english phonemes using signals from functional speech motor cortex. Journal of neural engineering, 11(3):035015, 2014.

[39] Klaus-Robert Müller, Matthias Krauledat, Guido Dornhege, Gabriel Curio, and Benjamin Blankertz. Machine learning techniques for brain-computer interfaces. Biomed. Tech, 49(1):11–22, 2004.

[40] Luis Fernando Nicolas-Alonso and Jaime Gomez-Gil. Brain computer interfaces, a review. Sensors, 12(2):1211–1279, 2012.

[41] Thomas J Oxley, Nicholas L Opie, Sam E John, Gil S Rind, Stephen M Ronayne, Tracey L Wheeler, Jack W Judy, Alan J McDonald, Anthony Dornom, Timothy JH Lovell, et al. Minimally invasive endovascular stent-electrode array for high-fidelity, chronic recordings of cortical neural activity. Nature biotechnology, 34(3):320–327, 2016.

[42] Miguel Pais-Vieira, Mikhail Lebedev, Carolina Kunicki, Jing Wang, and Miguel Nicolelis. A brain-to-brain interface for real-time sharing of sensorimotor information. Scientific reports, 3:1319, 02 2013.

[43] Artur Petrosyan, Mikhail Sinkin, Mikhail Lebedev, and Alexei Ossadtchi. Decoding and interpreting cortical signals with a compact convolutional neural network. Journal of Neural Engineering, 18(2):026019, 2021.

[44] Nick F Ramsey, Efraïm Salari, Erik J Aarnoutse, Mariska J Vansteensel, Martin G Bleichner, and ZV Freudenburg. Decoding spoken phonemes from sensorimotor cortex with high-density ecog grids. Neuroimage, 180:301–311, 2018.

[45] Philémon Roussel, Gaël Le Godais, Florent Bocquelet, Marie Palma, Jiang Hongjie, Shaomin Zhang, Anne-Lise Giraud, Pierre Mégevand, Kai Miller, Johannes Gehrig, et al. Observation and assessment of acoustic contamination of electrophysiological brain signals during speech production and sound perception. Journal of Neural Engineering, 17(5):056028, 2020.

[46] Philémon Roussel, Gaël Le Godais, Florent Bocquelet, Marie Palma, Jiang Hongjie, Shaomin Zhang, Philippe Kahane, Stéphan Chabardès, and Blaise Yvert. Acoustic contamination of electrophysiological brain signals during speech production and sound perception. BioRxiv, page 722207, 2019.

[47] Gerwin Schalk and Eric C Leuthardt. Brain-computer interfaces using electrocorticographic signals. IEEE reviews in biomedical engineering, 4:140–154, 2011.

[48] M Sinkin, A Osadchiy, M Lebedev, K Volkova, M Kondratova, I Trifonov, et al. High resolution passive speech mapping in dominant hemisphere glioma surgery. Russ. J. Neurosurg, 21:12–18, 2019.

[49] Frank Soong and B Juang. Line spectrum pair (lsp) and speech data compression. In ICASSP’84. IEEE International Conference on Acoustics, Speech, and Signal Processing, volume 9, pages 37–40. IEEE, 1984.

[50] Stanley Smith Stevens, John Volkmann, and Edwin Broomell Newman. A scale for the measurement of the psychological magnitude pitch. The journal of the acoustical society of america, 8(3):185–190, 1937.

[51] Pengfei Sun, Gopala K Anumanchipalli, and Edward F Chang. Brain2char: a deep architecture for decoding text from brain recordings. Journal of Neural Engineering, 17(6):066015, 2020.

[52] Christian Szegedy, Wei Liu, Yangqing Jia, Pierre Sermanet, Scott Reed, Dragomir Anguelov, Dumitru Erhan, Vincent Vanhoucke, and Andrew Rabinovich. Going deeper with convolutions. In Proceedings of the IEEE conference on computer vision and pattern recognition, pages 1–9, 2015.

[53] Ksenia Volkova, Mikhail A Lebedev, Alexander Kaplan, and Alexei Ossadtchi. Decoding movement from electrocorticographic activity: A review. Frontiers in neuroinformatics, 13:74, 2019.

[54] Sarah K Wandelt, Spencer Kellis, David A Bjånes, Kelsie Pejsa, Brian Lee, Charles Liu, and Richard A Andersen. Decoding grasp and speech signals from the cortical grasp circuit in a tetraplegic human. Neuron, 2022.

[55] Francis R Willett, Donald T Avansino, Leigh R Hochberg, Jaimie M Henderson, and Krishna V Shenoy. High-performance brain-to-text communication via handwriting. Nature, 593(7858):249–254, 2021.

[56] Guy H Wilson, Sergey D Stavisky, Francis R Willett, Donald T Avansino, Jessica N Kelemen, Leigh R Hochberg, Jaimie M Henderson, Shaul Druckmann, and Krishna V Shenoy. Decoding spoken english from intracortical electrode arrays in dorsal precentral gyrus. Journal of Neural Engineering, 17(6):066007, 2020.

[57] Min Xu, Ling-Yu Duan, Jianfei Cai, Liang-Tien Chia, Changsheng Xu, and Qi Tian. Hmm-based audio keyword generation. In Pacific-Rim Conference on Multimedia, pages 566–574. Springer, 2004.

